# Single-molecule long-read sequencing reveals the chromatin basis of gene expression

**DOI:** 10.1101/533158

**Authors:** Yunhao Wang, Anqi Wang, Zujun Liu, Andrew Thurman, Linda S. Powers, Meng Zou, Adam Hefel, Yunyi Li, Joseph Zabner, Kin Fai Au

## Abstract

Genome-wide chromatin accessibility and nucleosome occupancy profiles have been widely investigated, while the long-range dynamics remains poorly studied at the single-cell level. Here we present a new experimental approach MeSMLR-seq (methyltransferase treatment followed by single-molecule long-read sequencing) for long-range mapping of nucleosomes and chromatin accessibility at single DNA molecules, and thus achieve comprehensive-coverage characterization of the corresponding heterogeneity. We applied MeSMLR-seq to haploid yeast, where single DNA molecules represent single cells, and thus we could investigate the combinatorics of many (up to 356) nucleosomes at long range in single cells. We illustrated the differential organization principles of nucleosomes surrounding transcription start site for silently- and actively-transcribed genes, at the single-cell level and in the long-range scale. The heterogeneous patterns of chromatin statuses spanning multiple genes were phased. Together with single-cell RNA-seq data, we quantitatively revealed how chromatin accessibility correlated with gene transcription positively in a highly-heterogeneous scenario. Moreover, we quantified the openness of promoters and investigated the coupled chromatin changes of adjacent genes at single DNA molecules during transcription reprogramming.

## INTRODUCTION

In eukaryotic organisms, cells are faced with genetic information storage and packaging problems. As the carrier of genetic information, instead of folding into a disorganized yarn ball, DNA strands wrap around thousands of protein cores like “beads on a string”. As the fundamental unit of chromatin, nucleosome consists of ~147 bp DNA wrapping around a histone octamer composed of four core histones (H2A, H2B, H3 and H4) (1). Nucleosomes are connected by stretches of “linker DNA”. Dynamic packaging of nucleosomes results in two different chromatin accessibility statuses: open (accessible and active genomic regions with sparse nucleosome occupancy) and closed (inaccessible and inactive genomic regions with dense nucleosome occupancy). Positioning of nucleosomes and dynamic changes of chromatin status play important regulatory roles in DNA-templated processes such as transcription, DNA replication and repair (2).

Current genome-wide methods of nucleosome positioning and/or chromatin accessibility mapping are mainly based on three types of assays followed by short-read sequencing technologies: 1) nucleosome’s protection of nucleosomal DNA sequences from endogenous and exogenous enzymes (e.g., MNase-seq, DNase-seq, ATAC-seq, NOMe-seq and MPE-seq) (3–7); 2) chromatin immunoprecipitation using a specific histone antibody (e.g., ChIP-seq with H3) (8); and 3) solubility differences between nucleosomal DNA and naked linker DNA (e.g., FAIRE-seq) (9). In particular, NOMe-seq treats target sample with exogenous methyltransferase to detect nucleosome positioning and chromatin accessibility: the nucleosome protects nucleosomal DNA from being methylated by exogenous methyltransferase, while cytosines in naked linker DNA sequences are methylated to 5-methylcytosine (5mC) (6). The following bisulfite sequencing identifies this methylation profile as bisulfite can convert unmethylated cytosine to uracil, which discriminates 5mC from unmethylated cytosine.

These methods can map averaged patterns of nucleosome positioning and chromatin accessibility in a population of cells, failing in precise identification at the single-cell level. Although the single-cell versions of the methods have been recently developed (10–16), the corresponding sparse sequencing coverage and short read length lack information for addressing complex long-range chromatin status and nucleosome positioning. Therefore, the heterogeneity of nucleosome positioning and chromatin accessibility is rarely studied. Moreover, it is even more challenging to define nucleosome positioning patterns and dynamics and chromatin accessibility at single DNA molecules, so it is hard to detect subtle but meaningful differences between seemingly identical cells. This is a critical gap of understanding the mechanism of how nucleosomes assemble, disassemble and slide.

The emerging single-molecule long-read sequencing technology (i.e., Oxford Nanopore Technologies, ONT) provides unique data features that are possible to fill the gap: 1) 5mC can be directly detected at the single-base resolution at the single-molecule level based on ONT electrolytic current signal dynamics without bisulfite conversion (17, 18); 2) unlike the other sequencing platforms (such as Sanger sequencing and Second Generation Sequencing (SGS, e.g., Illumina)), PCR amplification is not required for ONT sequencing, so each ONT read can reveal the genomic events at the single-molecule level; 3) ONT reads are ultra-long (up to 2.3 Mb) (19) so that they can cover combinatorics of many nucleosomes and different chromatin statuses spanning multiple genomic elements. Leveraging the informative ONT sequencing technology, we developed an experimental approach MeSMLR-seq (methyltransferase treatment followed by ONT single-molecule long-read sequencing) and the corresponding bioinformatics method, so as to investigate heterogeneous and dynamic insight of long-range chromatin status and nucleosomes. Instead of bisulfite conversion (with PCR amplification) and short-read sequencing, the footprint of exogenous 5mCs from GpC-specific methyltransferase treatment is detected at single DNA molecules (without any PCR amplification) by ONT sequencing in the MeSMLR-seq protocol, and is next used to detect nucleosome positioning and chromatin accessibility computationally.

We applied MeSMLR-seq to haploid *Saccharomyces cerevisiae* cells, where single DNA molecules represent single cells, so it allows the “one-to-one” link between sequencing read (i.e., sequencing molecule) and haploid cell. Thus, each single MeSMLR-seq read can be used to mimic single cell in a given genomic region and the heterogeneity can be investigated without single-cell sequencing. We showed consistent and comparable bulk-level nucleosome occupancy profiles generated by MeSMLR-seq and MNase-seq, and demonstrated the accuracy and robustness of MeSMLR-seq on single-molecule long-range mapping of nucleosomes, and investigated the organization principle of nucleosomes surrounding transcription start site (TSS). Next, we evaluated the performance of MeSMLR-seq on chromatin accessibility mapping and showed the heterogeneity of combinatorial chromatin statuses over multiple genomic regions. In addition, with the unique MeSMLR-seq output, the relationship between chromatin accessibility and gene transcription was investigated quantitatively. Moreover, we revealed the coupled chromatin changes of adjacent genes during transcription reprogramming.

## RESULTS

### Overview of MeSMLR-seq

In brief, the experimental approach MeSMLR-seq (**Me**thyltransferase treatment followed by **S**ingle-**M**olecule **L**ong-**R**ead sequencing) contains two main steps: 1) methyltransferase (M.CviPI) treatment to convert cytosine to 5mC at GpC sites at naked linker DNA and open chromatin; and 2) ONT sequencing to detect 5mC profile that is subsequently used to identify nucleosome positioning and chromatin accessibility (Fig. 1). The first step has been shown feasible at both bulk-cell and single-cell level by NOMe-seq and the other previous studies (13–16). In addition, ONT has been reported to detect 5mC at CpG sites (17, 18), based on which an in-house tool was developed to map 5mC profile at GpC sites for MeSMLR-seq data (see “Nucleosome positioning detection at the single-molecule level”).

**Fig. 1.**
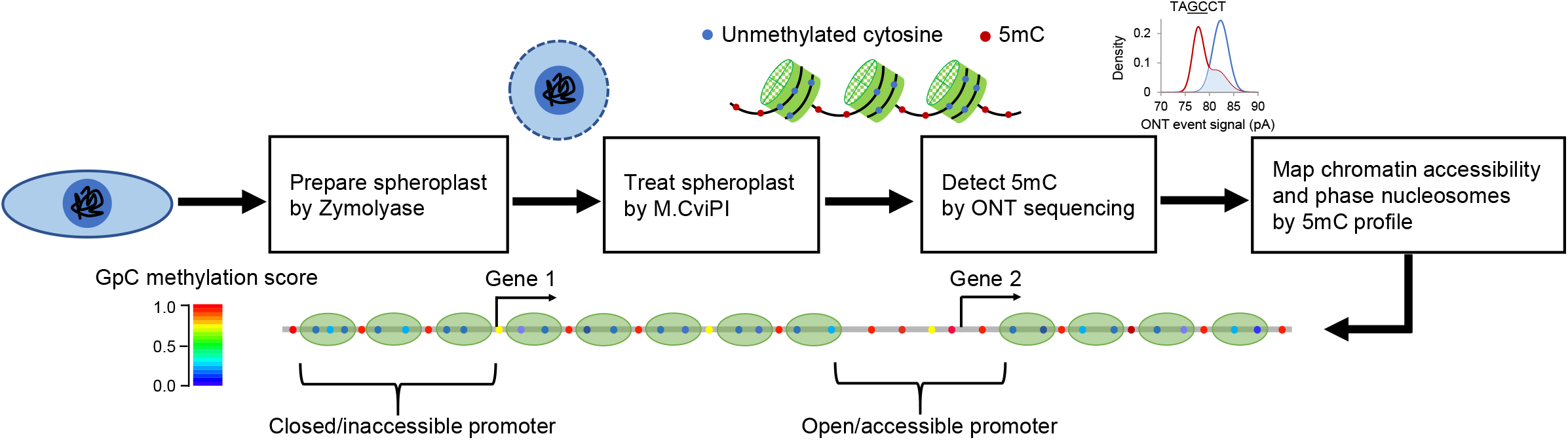
Overview of MeSMLR-seq. Experimental approach (methyltransferase treatment plus ONT sequencing) in yeast and the corresponding bioinformatics analyses (5mC detection, chromatin accessibility mapping and nucleosome phasing).

In the proof-of-concept application of MeSMLR-seq to haploid *Saccharomyces cerevisiae* (BY4741 strain), an additional step was applied to digest cell wall that serves as a barrier against methyltransferase treatment to genomic DNA: yeast cells were treated with Zymolyase to generate spheroplasts (Fig. 1 and SI Appendix, Fig. S1). After the subsequent methyltransferase treatment, extracted genomic DNA without any PCR amplification was directly submitted to library preparation and ONT sequencing. The genomic DNA that undergoes *in vivo* spheroplast methylation was referred as target sample of MeSMLR-seq. In addition, we prepared negative control and positive control samples as training data for 5mC detection (see the below section “Nucleosome positioning detection at the single-molecule level”, and SI Appendix, Fig. S1). Native genomic DNA extracted from yeast without M.CviPI treatment was used as negative control (all cytosines at GpC sites were unmethylated) since there is no endogenous 5mC on yeast genome as previously reported (20). Genomic DNA treated by M.CviPI (without spheroplast methylation) was used as positive control (all cytosines at GpC sites were converted to 5mCs).

As the efficiency of M.CviPI methylation served a critical role in the whole protocol, it was evaluated at selected genomic regions by bisulfite sequencing as previously described (16). The methylation efficiency of positive control sample was 99.37% and 13 single colonies of the selected region from target sample were all successfully-methylated, indicating the high methylation efficiency.

Using ONT GridION platform with R9.4.1 chemistry, we sequenced one flow cell per sample and generated 0.9 million (positive control), 1.2 million (negative control) and 1.3 million (on average for six target samples) reads (i.e., sequencing molecules), separately, which were uniquely aligned to yeast genome (SI Appendix, Table S1). The longest sequencing molecule was 63.1 kb. In particular, from the target sample where yeast was grown in rich media (1% yeast extract, 2% peptone and 2% glucose) we generated 1.4 million sequencing molecules with the median length of 7.2 kb, covering 821X of yeast genome.

### Detection and phasing of nucleosome positioning at single DNA molecules

We first identified 5mC methylation status for every GpC site on each DNA molecule based on the ONT sequencing current signal (referred as event level). Since the previous studies (17, 18) showed the event level depended on the context sequence (e.g., 6-mer), our positive and negative control data were used to train signal distributions for each 6-mer containing target GpC dinucleotide under the occasions of methylation and unmethylation. The event levels of a given 6-mer from the target sample were compared with the corresponding trained distributions to obtain a posterior of methylation for every GpC site on each molecule, which we denoted as the methylation score (SI Appendix, Fig. S2A). There was no obvious bias of 5mC methylation calling between the molecules that were aligned to forward and reverse strands, and the areas under the receiver operating characteristic curve (AUC) were both 0.86 (Fig. 2A). Correlation analysis of methylation status of paired GpC sites at single molecules showed a remarkable pattern with period distance of 170-180 bp, which was the same as the length of nucleosomal DNA (147 bp) plus regular linker DNA (20-30 bp) (Fig. 2B). Therefore, we can identify nucleosome positioning at single molecules from the methylation profiles by developing the bioinformatics method NP-SMLR (Nucleosome Positioning detection by Single-Molecule Long-Read sequencing) as below.

**Fig. 2.**
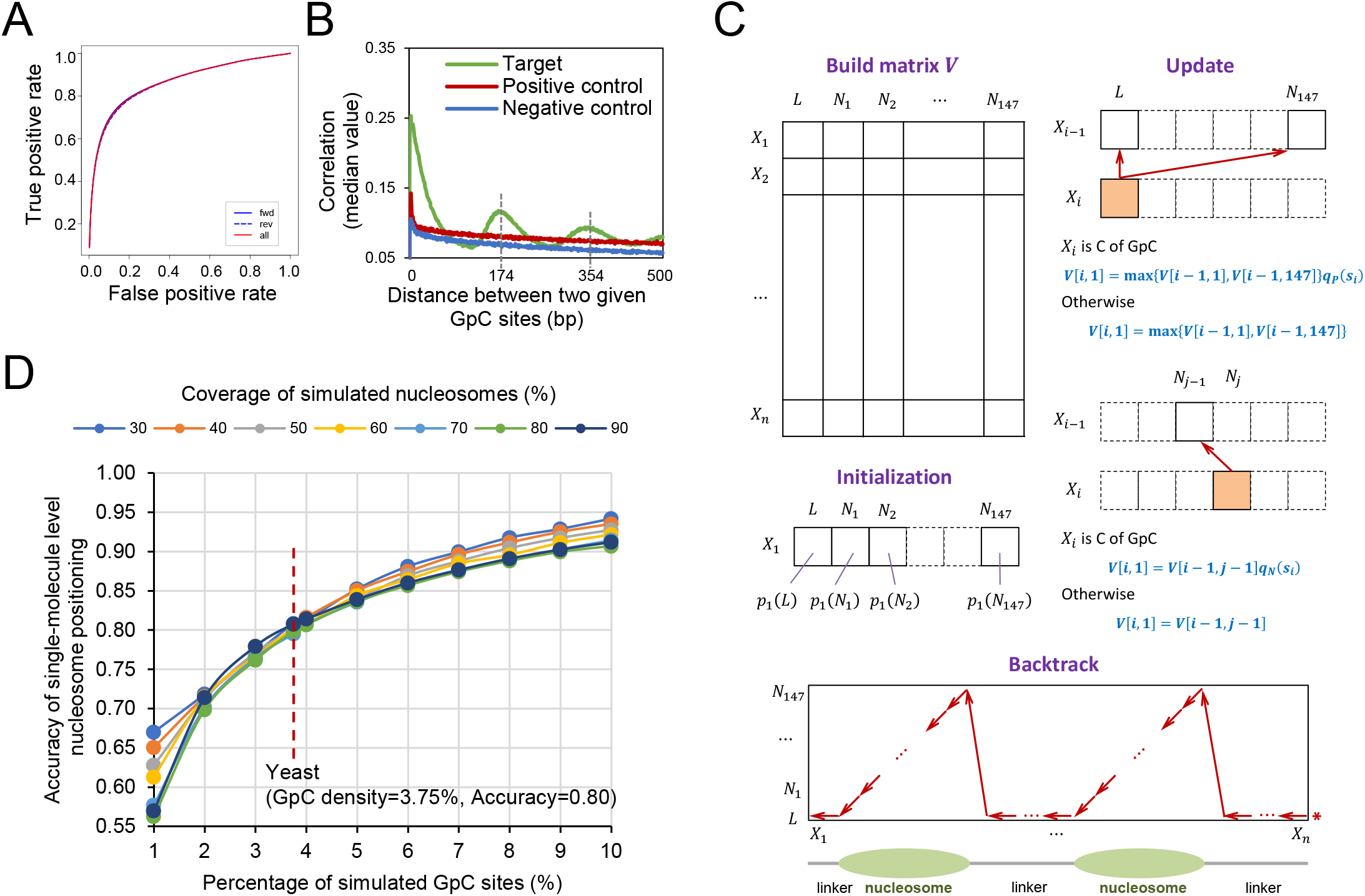
5mC detection and nucleosome positioning by MeSMLR-seq data. **A.** ROC curve of 5mC detection on GpC sites. The molecules that were aligned to forward (fwd) and reverse (rev) genomic strands were analyzed separately. **B.** Correlation coefficients between methylation scores of mutually paired GpC sites from the same molecules with respect to their corresponding distances. **C.** Dynamic programming algorithm for nucleosome positioning detection (NP-SMLR). A matrix regarding the nucleotide sequence (row) and nucleosomal statuses (column) is made, followed by initialization, iterative update for entries, and backtrack search for optimal path (see Materials and Methods for details). **D.** Accuracy of nucleosome positioning under different nucleosome coverage and GpC frequencies.

Let *X*_1_*X*_2_ … *X_l_* be a molecule, where *X*_*i*_ is the *i*-th base. Denote *s_i_* as the methylation score of *X_i_*, if *X_i_* is the cytosine of the GpC dinucleotide. Suppose that the methylation scores of all GpC sites are independent. Nucleosome positioning detection refers to finding a path *π* = *π*_1_*π*_2_ … *π_l_* that maximizes the likelihood of signals:

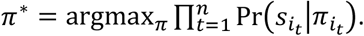

*π_i_* takes the value from {*L, N*_1_, *N*_2_, …, *N*_147_}. *L* represents the linker region; *N_m_* represents the *m*-th base within a nucleosome; *i*_1_, *i*_2_, …, *i_n_* are the positions of cytosines that belong to GpC dinucleotides. The elements of path *π* are restricted that: 1) *N_m_* is followed by *N_m+t_* (1 ≤ *m* ≤ 146); 2) *N*_147_ is followed by *L*; and 3) *L* is followed by *L* or *N*_1_. The problem is essentially an alignment between a sequence of nucleotides and a sequence of nucleosomal statuses. NP-SMLR adopts dynamic programming algorithm (21) for solution: a matrix regarding the nucleotide sequence and nucleosomal statuses is made, entries are updated iteratively, and the optimal path is obtained through backtracking (Fig. 2C and SI Appendix, Fig. S2B).

Due to the lack of more advanced experimental technology to generate gold standard, we evaluated the accuracy of nucleosome positioning detection at the single-molecule level by simulation tests. The tests were performed under different settings of nucleosome coverage (proportion of bases covered by nucleosomes, ranges from 30-90%) and GpC frequency (ranges from 1-10%) (Fig. 2D). The accuracy increased with GpC frequency, while the effect of nucleosome coverage was mild. In case of yeast genome with 3.75% density of GpC sites, NP-SMLR was very robust to reach the accuracy of 80% regardless different nucleosome coverage, which represented different scenarios of chromatin status (Fig. 2D).

### Performance of nucleosome positioning detection at the bulk-cell level

In terms of nucleosome positioning at the bulk-cell level, MeSMLR-seq provided comparable results with the widely-used method MNase-seq (SI Appendix, section 1 and section 2) (22, 23). The averaged Pearson’s correlation coefficient between three MeSMLR-seq data (forwardly, reversely aligned molecules and their combination) and three MNase-seq replicates was 0.75 (Fig. 3A). 77% nucleosomes called by MeSMLR-seq were also detected by MNase-seq (Fig. 3C). For example of the *DAL* (degradation of allantoin) gene cluster, the nucleosome peaks called by MeSMLR-seq and MNase-seq were generally well aligned (Fig. 3B). In long-range scale, single MeSMLR-seq reads can phase a number of nucleosomes (median number was 37 and maximal number was 356 in our data), so it captures the dynamics and heterogeneity of nucleosome positioning among DNA molecules (Fig. 3D and SI Appendix, Table S2). For instance, 35 to 61 nucleosomes (median number 58) were phased at the single molecules covering the *DAL* gene cluster across a 10 kb genomic region (Fig. 3E), which illustrated large-range variation as well as local subtle difference of nucleosome positioning.

**Fig. 3.**
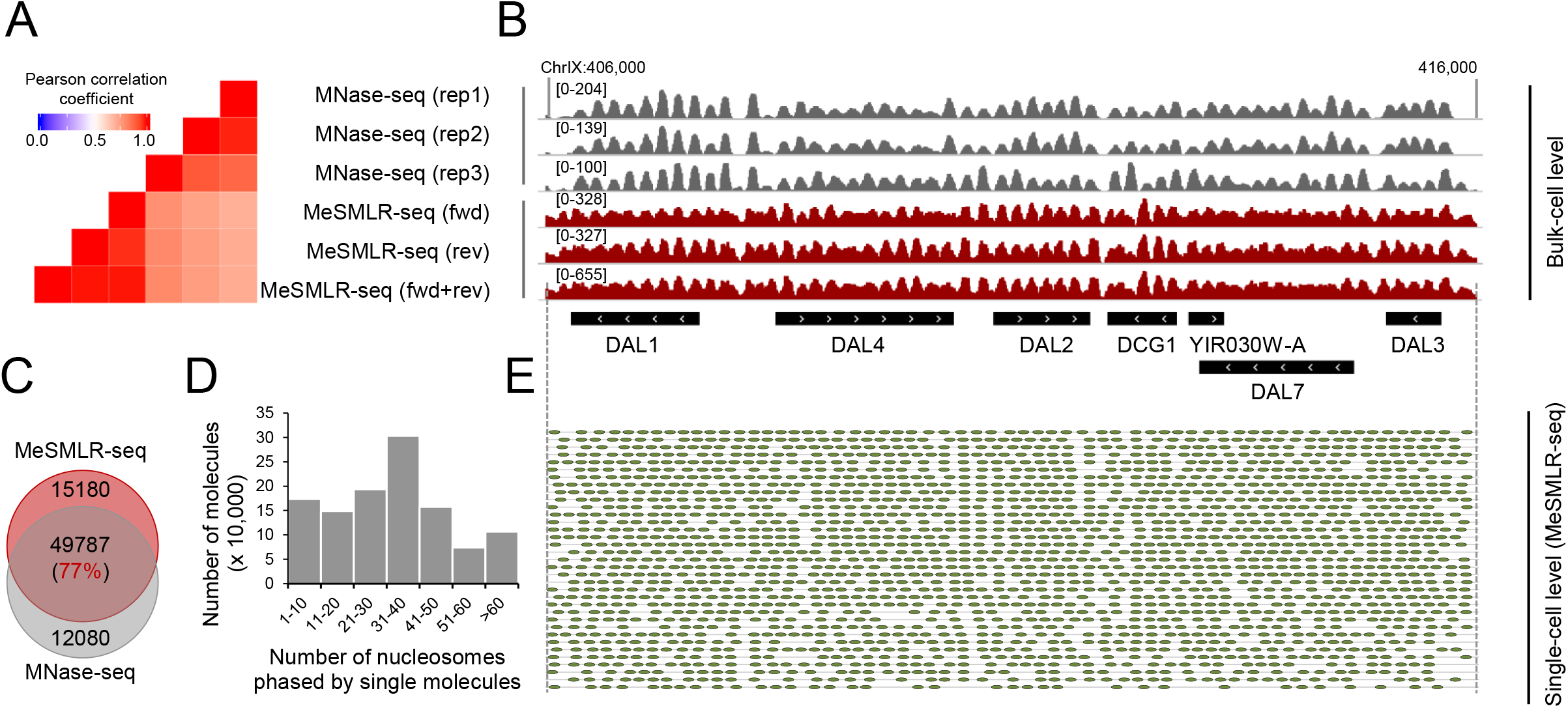
Performance evaluation of MeSMLR-seq on bulk-level nucleosome occupancy, and single-molecule long-range phasing of nucleosomes. **A.** Correlation of nucleosome occupancy profiles generated by MeSMLR-seq and MNase-seq. For MeSMLR-seq, the molecules that were aligned to forward (fwd) and reverse (rev) genomic strands were analyzed separately. **B.** Nucleosome occupancy profiles at the bulk-cell level provided by MeSMLR-seq and MNase-seq. **C.** Overlap of nucleosomes detected by MeSMLR-seq and MNase-seq at the bulk-cell level. **D**. Number of nucleosomes phased at single sequencing molecules of MeSMLR-seq data under 2% glucose condition. **E.** Detection and phasing of nucleosomes at the single-molecule level by NP-SMLR. Each grey line represents a molecule. Green oval represents nucleosome.

### Direct long-range evidence of differential nucleosome organization

A few single-cell epigenome sequencing approaches have revealed the heterogeneity of chromatin status and nucleosome positioning within a cell population (10–16). Notably, Lai *et al*. recently reported the differential nucleosome organization principles for silent and active genes using single-cell MNase-seq (12) (Fig. 4A). However, these studies lacked a long-scale nucleosome positioning scene at the single-cell resolution due to short sequencing length and sparse data coverage within single cells. As shown above, MeSMLR-seq can determine the heterogeneous long-range phasing of nucleosomes, so we can investigate nucleosome organization logic in a comprehensive way (Fig. 3E).

**Fig. 4.**
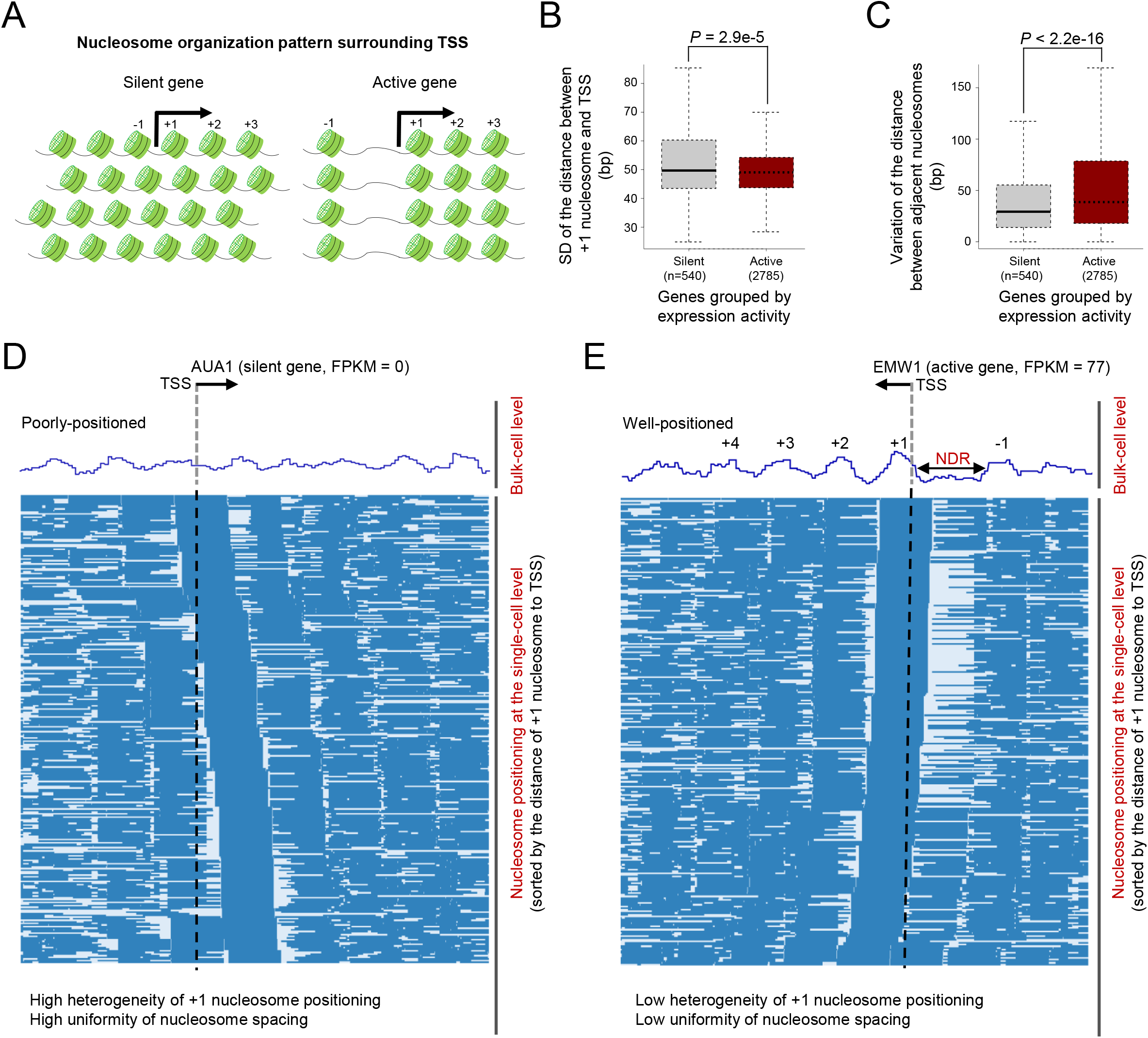
Differential nucleosome organization principles for silent and active genes. **A.** Previous studies revealed nucleosome organization patterns surrounding TSS of silent (left) and active (right) genes (12). Nucleosome positioning in promoter regions of silent genes showed large variation among cells but was highly uniformly spaced within each cell. In contrast, nucleosome positioning surrounding TSS of active genes showed little variation among cells but relatively non-uniformly spacing within each cell. **B.** Heterogeneity of nucleosome positioning for silent (FPKM=0) and active (FPKM>50) genes. The heterogeneity of nucleosome positioning was measured by the standard deviation (SD) of the distances between +1 nucleosomes and TSS. The p-value was calculated by Wilcoxon rank sum test. **C.** Uniformity of nucleosome spacing for silent (FPKM=0) and active (FPKM>50) genes. See Materials and Methods for the definition of uniformity. The p-value was calculated by Wilcoxon rank sum test. **D.** Long-range nucleosome positioning patterns for the silently-transcribed gene *AUA1* across different cells. Each row represents a cell and nucleosome is labeled as blue bar. **E.** Long-range nucleosome positioning patterns for the actively-transcribed gene *EWM1* across different cells.

We focused on the nucleosome organization surrounding TSS, which plays important role in transcription regulation (24). For each gene, we measured the heterogeneity of nucleosome positioning by the standard deviation of the distances between +1 nucleosome and TSS over all single cells. Compared to active genes, silent genes showed larger heterogeneity of nucleosome positioning among different cells (Fig. 4B and SI Appendix, Fig. S3A). Next, we evaluated the uniformity of nucleosome spacing within single cells by the variation of the distance between adjacent nucleosomes. In contrast to active genes, the nucleosomes surrounding TSS of silent genes were more uniformly spaced (Fig. 4C and SI Appendix, Fig. S3B). For instance, at the bulk-cell level, nucleosomes surrounding TSS of the lowly-expressed gene *AUA1* (FPKM=0) were poorly positioned (Fig. 4D), while there were well-positioned nucleosomes (including −1, +1, +2, +3 and +4 nucleosomes) surrounding TSS of the active gene *EMW1* (FPKM=77) and a pronounced nucleosome-depletion region (NDR) in the upstream of TSS (Fig. 4E). At the single-cell level, the positioning of +1 nucleosome of *AUA1* had a remarkable continuous shift pattern across different cells, whereas it was relatively steady for *EMW1* (Fig. 4D, E). Compared with *EMW1*, the distances between +1 nucleosomes and TSS for *AUA1* were more approximate to a uniform distribution (SI Appendix, Fig. S4A, B), which represented the ideal occasion for continuous shift pattern. In addition, the spacing of nucleosomes surrounding of TSS of *AUA1* was relatively uniform within single cells (Fig. 4D and SI Appendix, Fig. S4C), while there was a pronounced NDR in the upstream of TSS of *EWM1*, which disrupted the uniformity of nucleosome spacing (Fig. 4E and SI Appendix, Fig. S4D). MeSMLR-seq resolves these differential nucleosome organization principles with direct and convincing evidence at a long-range scale from single molecules/cells that are hard to be obtained by the bulk-cell and short-read sequencing approaches.

### Single-molecule long-range measurement of chromatin accessibility

Based on the methylation profiles of MeSMLR-seq data, we also mapped the chromatin accessibility of yeast genome at both bulk-cell level and single-molecule level. To assess the performance on the bulk-cell chromatin accessibility mapping, we compared MeSMLR-seq with two widely-used methods, ATAC-seq (25) and DNase-seq (26) (SI Appendix, section 1 and section 3). Genome-wide chromatin accessibility profile revealed by MeSMLR-seq data was highly consistent with ATAC-seq (averaged Pearson’s r=0.80) and DNase-seq (averaged Pearson’s r=0.82) (Fig. 5A, B and SI Appendix, Fig. S5). In addition, >83% (1,615/1,934) significantly-accessible regions called by MeSMLR-seq were also supported by either ATAC-seq or DNase-seq (Fig. 5C). These results indicate that MeSMLR-seq provides comparable results with the existing methods on the bulk-cell level chromatin accessibility mapping.

**Fig. 5.**
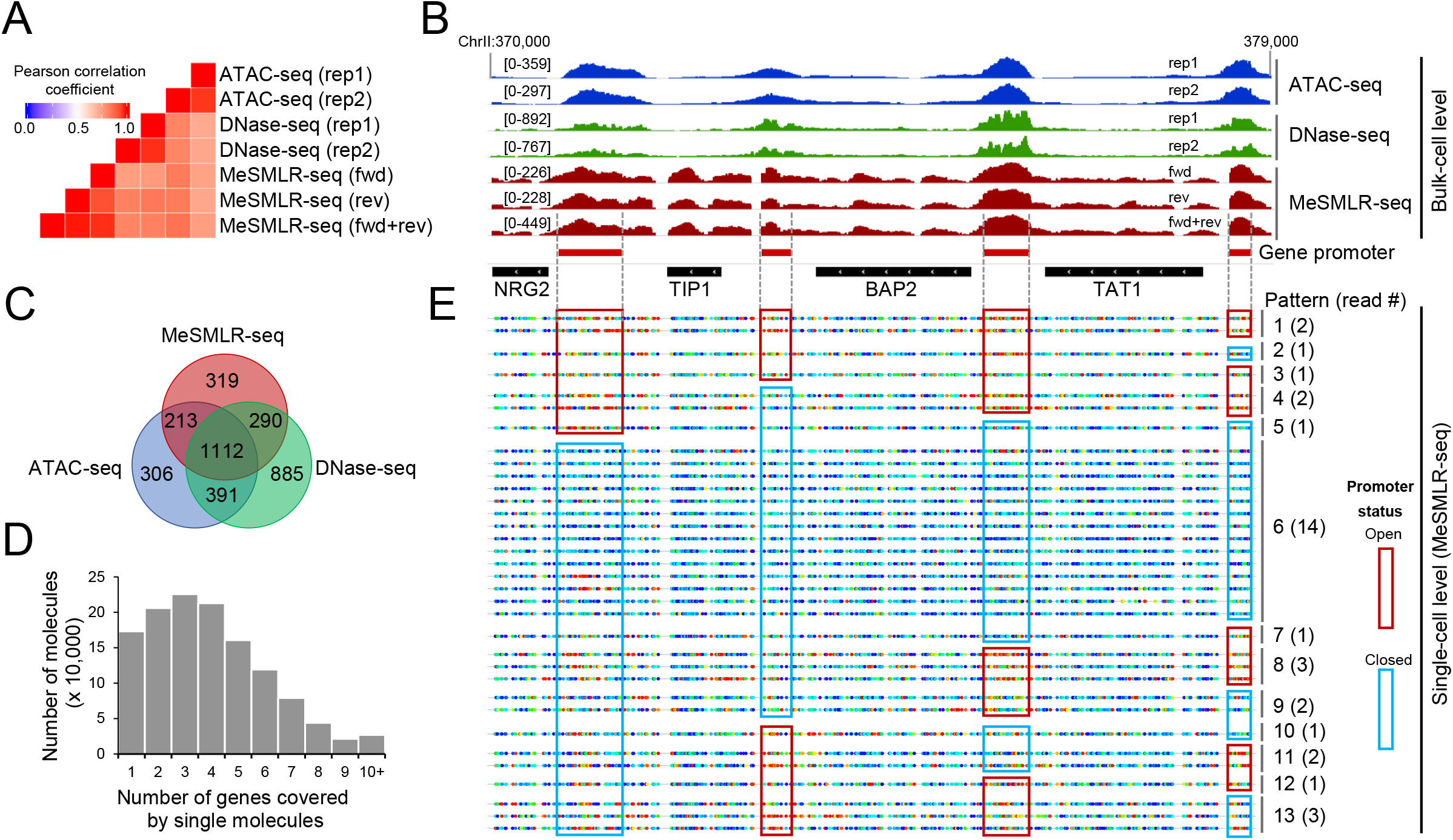
Performance evaluation of MeSMLR-seq on bulk-level chromatin accessibility mapping, and single-molecule long-range mapping of chromatin accessibility. **A.** Correlation of chromatin accessibility profiles generated by MeSMLR-seq, ATAC-seq and DNase-seq. **B.** Chromatin accessibility profiles at the bulk-cell level provided by MeSMLR-seq, ATAC-seq and DNase-seq. **C.** Overlap of the significantly-accessible regions (peaks) called by MeSMLR-seq, ATAC-seq and DNase-seq. **D.** Number of genes covered by single sequencing molecules of MeSMLR-seq data under 2% glucose condition. **E.** Single-molecule long-range mapping of chromatin accessibility by MeSMLR-seq. Each line represents a molecule. GpC site is labeled as rainbow-color dot, with methylation score from 0 (blue) to 1.0 (red). Thirteen combinatorial patterns of the promoter status of four genes are shown with different numbers of supporting sequencing molecules/cells. A promoter was defined as “open” (highlighted by red box) if the methylation scores of the including GpC sites had a median value greater than 0.5, and “closed” (highlighted by blue box) otherwise.

At the single-molecule level, a MeSMLR-seq read can fully cover multiple adjacent genes (median number was 4 and maximal number was 40 in our data), therefore we could examine the long-range chromatin accessibility at the single-molecule/-cell level (Fig. 5D and SI Appendix, Table S3). For example, 34 MeSMLR-seq molecules fully covered the 9 kb genomic region ChrII:370000-379000 that encompasses four genes (*NRG2, TIP2, BAP2* and *TAT1*). Based on the 5mC footprint, we identified the chromatin status (“open” or “closed”) of the promoters for four genes on each molecule and thus defined and quantified the coupled chromatin status patterns. In total, these molecules detected 13 out of 16 (4^2^, four genes with binary status “open” or “closed”) possible combinatorial patterns of the coupled chromatin statuses of four gene promoters (Fig. 5E). For instance, four genes in Pattern 1 (supported by 2 molecules) all had “open” promoters, whereas the promoters of four genes were all closed in Pattern 6 (supported by 14 molecules). Therefore, MeSMLR-seq is applicable to analyze the coupled chromatin statuses of adjacent genes and to investigate the heterogeneity of chromatin status within a cell population, which is challenging for the existing methods.

### Heterogeneous openness of gene promoter

Leveraging the single-molecule and long-range information of MeSMLR-seq data, we can discover and measure different levels of promoter openness instead of binary status. In the promoter region (ChrXVI:66400-67550) of the cell cycle regulation gene *CLN2*, the bulk-level chromatin accessibility profiles generated by the existing methods and MeSMLR-seq all showed a significant openness (Fig. 6A), while it was not clear if the promoters of *CLN2* among all cells were open, or if the open regions were similar in size. Based on the single-molecule nucleosome positioning profiles in the promoter region, 304 molecules that fully covered this region were clustered into three groups with different levels of promoter openness: closed (Cluster 1 with 176 molecules), narrowly-open (Cluster 2 with 75 molecules) and widely-open (Cluster 3 with 53 molecules) (Fig. 6B, right panel). The 5mC profiles at the molecules from three clusters also showed the remarkable difference of the widths of openness (Fig. 6B, left panel). This unique output of MeSMLR-seq is bringing new opportunities to perform quantitative analysis of the heterogeneous and dynamic promoter status.

**Fig. 6.**
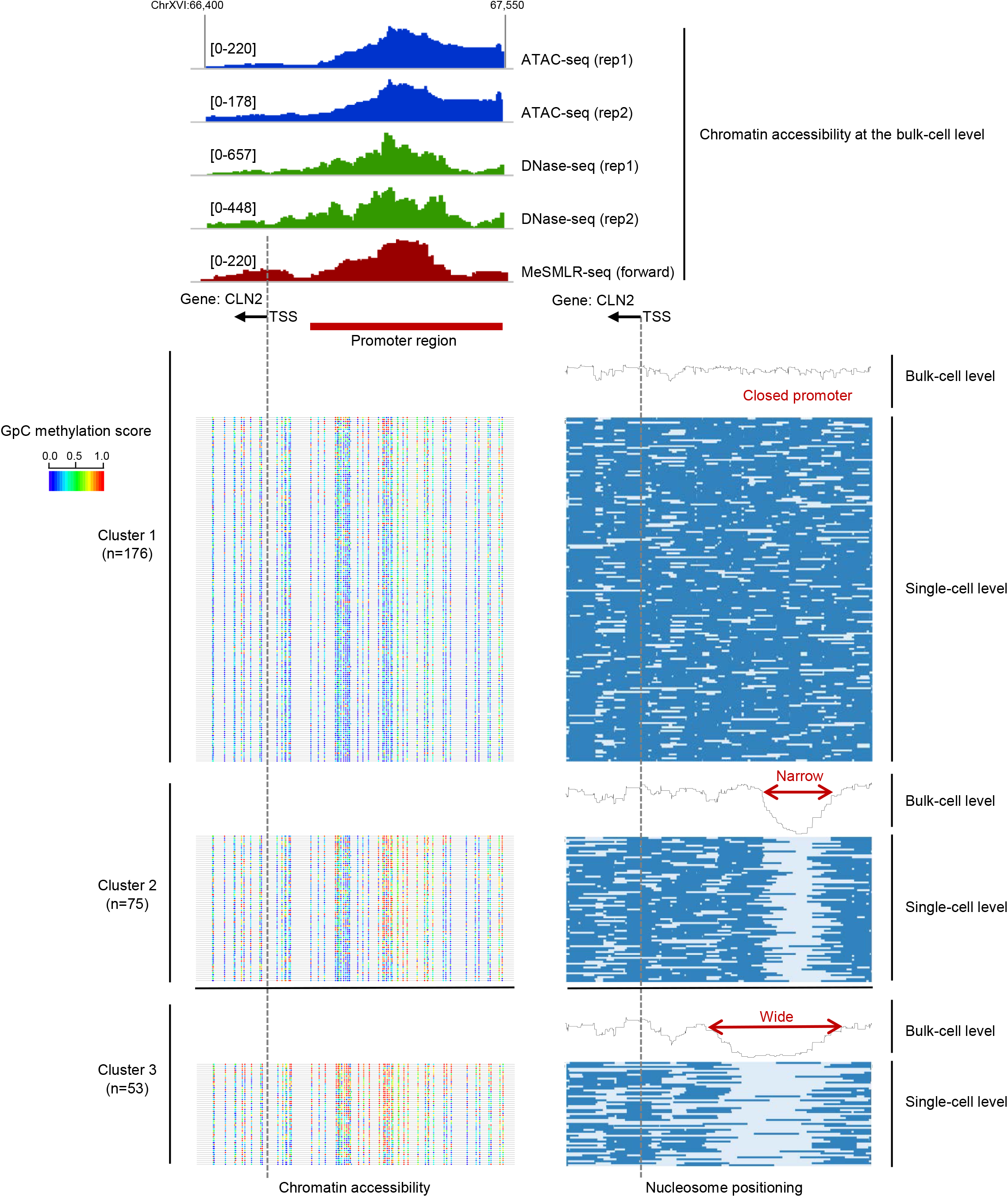
Heterogeneous promoter openness of *CLN2* in a cell population revealed by MeSMLR-seq. The bulk-level chromatin accessibility profiles (upper panel) were provided by ATAC-seq, DNase-seq, and MeSMLR-seq. MeSMLR-seq molecules were clustered into three groups with different promoter openness (by *k*-means clustering of the nucleosome positioning profiles, bottom right panel): closed, narrow open and wide open. Each row represents a molecule (i.e., a cell) and nucleosome is labeled as blue bar. The corresponding methylation profiles at GpC sites on each molecule are shown on the bottom left panel. Each line represents a molecule (i.e., a cell). GpC site is labeled as rainbow-color dot, with methylation score from 0 (blue) to 1.0 (red).

### Promoter openness and gene transcription

Using the MeSMLR-seq data, we generated the nucleosome occupancy profiles surrounding the TSSs of all protein-coding genes. Consistent with previous studies (22, 27), MeSMLR-seq data showed that highly-expressed genes had more pronounced nucleosome-depletion region in the upstream of TSS and well-positioned nucleosome array across gene body (Fig. 7A, B). Nucleosome occupancy of the genes with high expression levels showed an obvious drop at TSS and distinct peaks within gene body, while such tendency was mild for the genes with the lower 25^th^ percentile expression level (Fig. 7B).

**Fig. 7.**
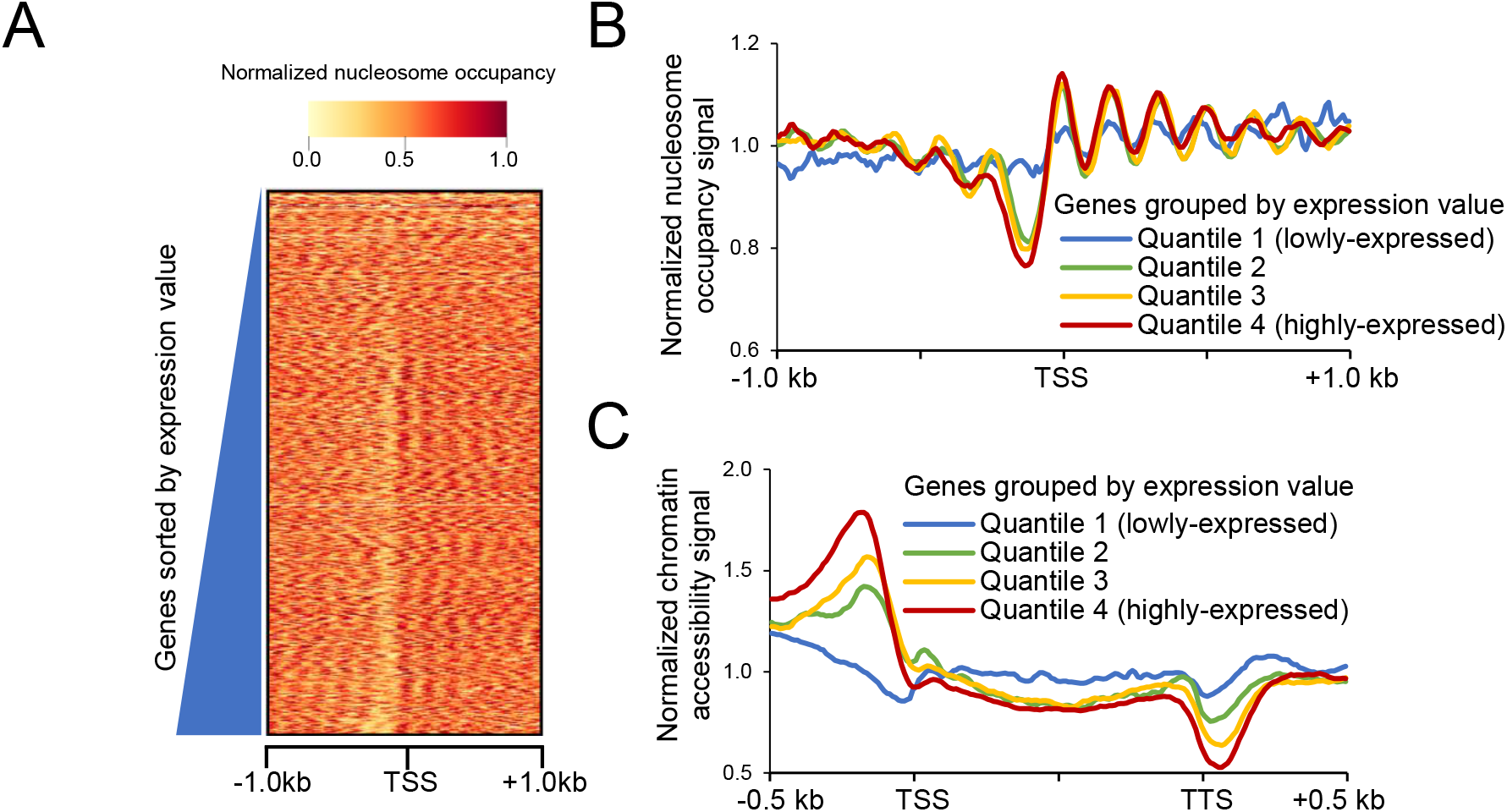
Relationship between nucleosome occupancy, chromatin accessibility and gene expression. **A.** Nucleosome occupancy profiles across all protein-coding genes with the ascending order of gene expression level from top to bottom. **B.** Nucleosome occupancy profiles at the bulk-cell level for protein-coding genes with different expression levels. **C.** Chromatin accessibility profiles at the bulk-cell level for protein-coding genes with different expression levels.

In addition to nucleosome occupancy, the chromatin accessibility profiles by MeSMLR-seq showed that the promoter regions of the highly-expressed genes were more accessible than the lowly-expressed genes (Fig. 7C). It indicates the critical role of promoter accessibility on gene transcription regulation. We further examined the chromatin statuses of the binding regions of several important transcriptional regulators, including RNA polymerase II (Pol2), five general regulatory factors (Abf1, Cbf1, Mcm1, Rap1 and Reb1) and two mediators (Med8 and Med17) (SI Appendix, section 1) (28–30). The enrichment signal of Pol2 in gene body was positively correlated with chromatin accessibility of gene promoter (SI Appendix, Fig. S6A). The binding regions of the other regulatory factors and mediators were relatively accessible and nucleosome-evicted, which allows the assembly of transcription initiation complex (SI Appendix, Fig. S6B-E).

### Dynamic change of chromatin status in response to different carbon sources

We next sought to investigate the dynamics of chromatin status during transcription changes in response to different nutrition conditions. Carbon source is the basic nutrition and is essential for yeast growth (31). In addition to glucose (Glu), which is the preferred carbon source for *S. cerevisiae*, we grew yeast cells separately using galactose (Gal) and raffinose (Raf) carbon sources, and generated both MeSMLR-seq and RNA-seq data. Compared with those under Gal and Raf conditions, yeast cells under Glu showed more accessible promoter (Fig. 8A). 21.62% (1,384 of 6,713) of protein-coding genes were differentially expressed between Glu and Gal, and 20% (1,332 of 6,713) between Glu and Raf, which indicated significant transcription reprogramming in response to different carbon sources (Fig. 8B). The up-regulated genes in Glu compared to Gal or Raf were mainly located in cytoplasm and involved in the biogenesis of ribosomes (Fig. 8C). In contrast, the up-regulated genes in both Gal and Raf conditions compared to Glu were significantly related to oxidation-reduction process, carbon metabolism, and located in mitochondrion. Those significantly up-regulated genes in Glu underwent more remarkable difference of chromatin accessibility in their promoters (*p*-value=1.2e-14 for Glu vs. Gal, *p*-value=3.6e-11 for Glu vs. Raf, Wilcoxon rank sum test, Fig. 8D), which contributed the overall high chromatin accessibility in preferred carbon source (Glu) over Gal and Raf (Fig. 8A).

**Fig. 8.**
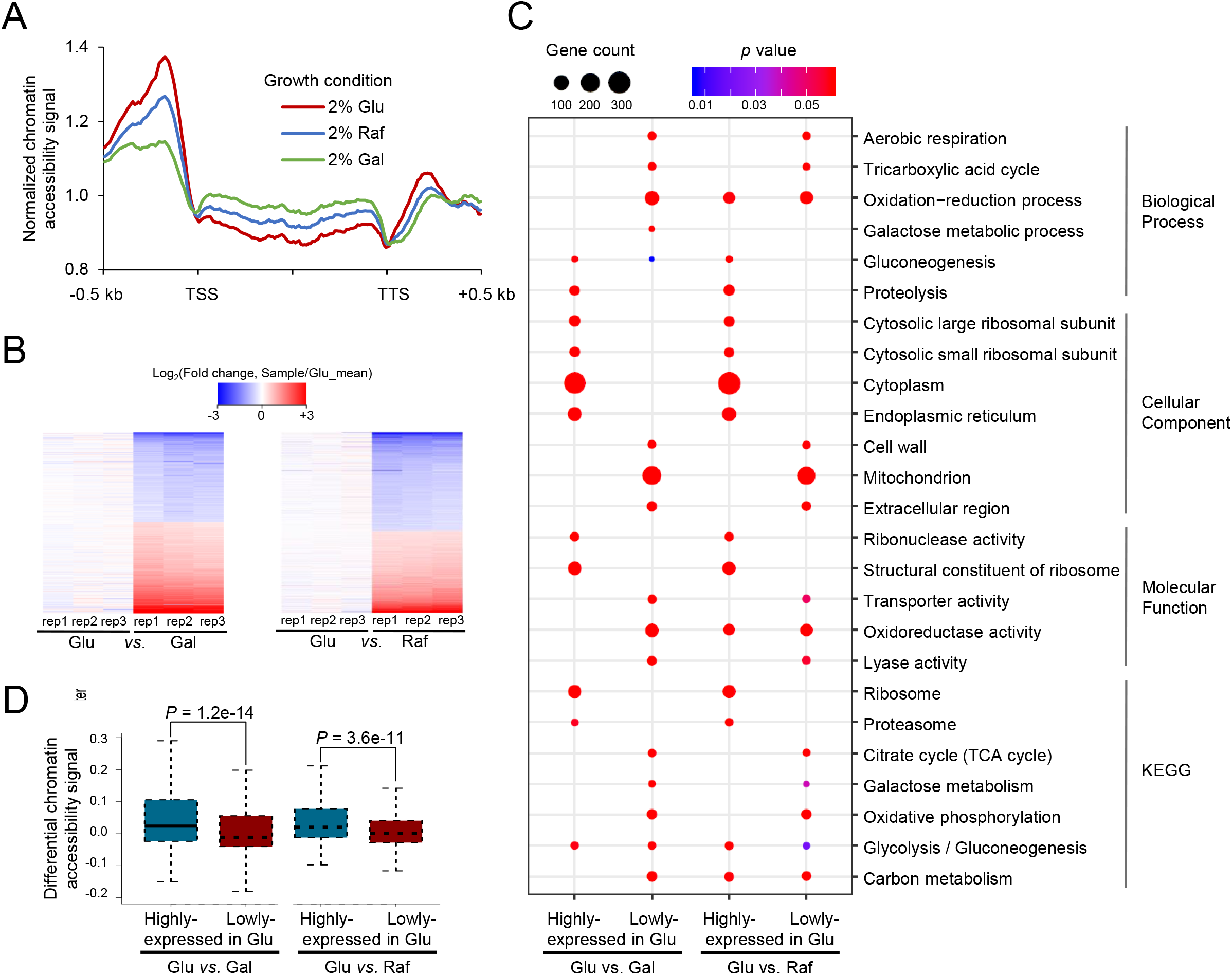
Differential chromatin accessibility and gene expression under different carbon sources. **A.** Differential chromatin accessibility patterns under glucose, galactose and raffinose. **B.** Differential gene expression patterns under different growth conditions. Fold change = (the FPKM value of the sample)/(the averaged FPKM under glucose condition). **C.** Gene enrichment analyses for differentially-expressed genes. **D.** Difference of chromatin accessibility between up- and down-regulated genes under different carbon sources. Glu, glucose; Gal, galactose; and Raf, raffinose.

### Quantitative relationship between gene expression and chromatin accessibility in cell population

Though the analyses above showed that the promoters of the highly-expressed genes over a cell population were generally more accessible than the low-expressed genes (Fig. 7), the quantitative relationship between promoter openness and gene transcription in a cell population remained unclear. Based on unique MeSMLR-seq data, we were able to calculate the fraction of cell subpopulation with open promoter of a given gene. With single-cell RNA-seq data for 2,812 yeast cells generated in this study (SI Appendix, section 4), we also calculated the fraction of cells with expression (read count ≥1) of a given gene (referred as expression frequency). The expression frequency within a cell population was positively correlated with the fraction of cells with open promoter (Fig. 9A). For example, the genes with open promoter in ≥40% cells had significantly larger expression frequency than the ones with open promoter in <10% cells (p-value <2.2e-16, Wilcoxon rank sum test, Fig. 9A). When grouping the genes based on expression frequency, we observed similar positive correlation (Fig. 9B). In addition, considering the bulk-cell expression, the highly-expressed genes had relatively large fractions of cell subpopulation with open promoter in comparison to the lowly-expressed ones (p-value <2.2e-16, Wilcoxon rank sum test, Fig. 9C). These results suggest that chromatin accessibility of promoter at the single-molecule/-cell level detected by MeSMLR-seq data can contribute to the prediction of gene expression level and frequency in a cell population.

**Fig. 9.**
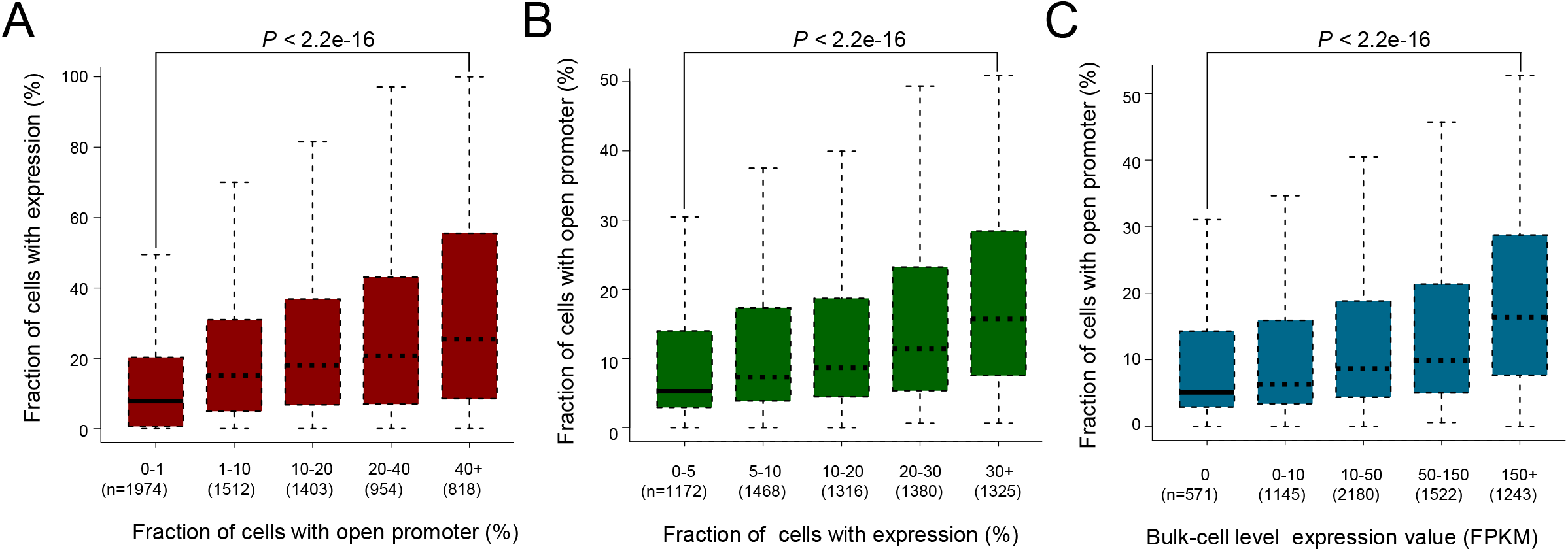
Quantitative relationship between chromatin accessibility and gene expression. **A, B.** Quantitative relationship between chromatin accessibility and gene expression in a cell population. The former was measured by the fraction of cells with open promoter, and the latter by the fraction of cells with expression (based on single-cell RNA-seq data). Genes were binned by one of the indices and the distribution of the other is shown. The gene was considered as “expressed” in a cell if the corresponding UMI (unique molecular identifier) count was ≥1. **C.** Quantitative relationship between the bulk-cell gene expression and the cell population ratio of open promoter. Genes were binned based on the bulk-cell gene expression level (RNA-seq data).

### Coupled chromatin accessibility changes of adjacent genes during transcription reprogramming

Making full use of the single-molecule and long-range advantages of MeSMLR-seq data, we explored the coupled chromatin status changes of two adjacent glucose transporter genes, *HXT3* and *HXT6* during transcription reprogramming. The transport of glucose across the plasma membrane is the first step of glucose metabolism, and the glucose (also called hexose) transporter genes play essential regulatory roles in glucose sensing, signaling and utilization in a yeast cell (32). HXT3 and HXT6 have different affinities to glucose (low-affinity for HXT3 and high-affinity for HXT6) and thus respond differently to the change of glucose concentration. With the decrease of glucose concentration, the expression of *HXT3* decreased whereas *HXT6* increased, which corresponded to their low- and high-affinity of glucose (Fig. 10A, B).

**Fig. 10.**
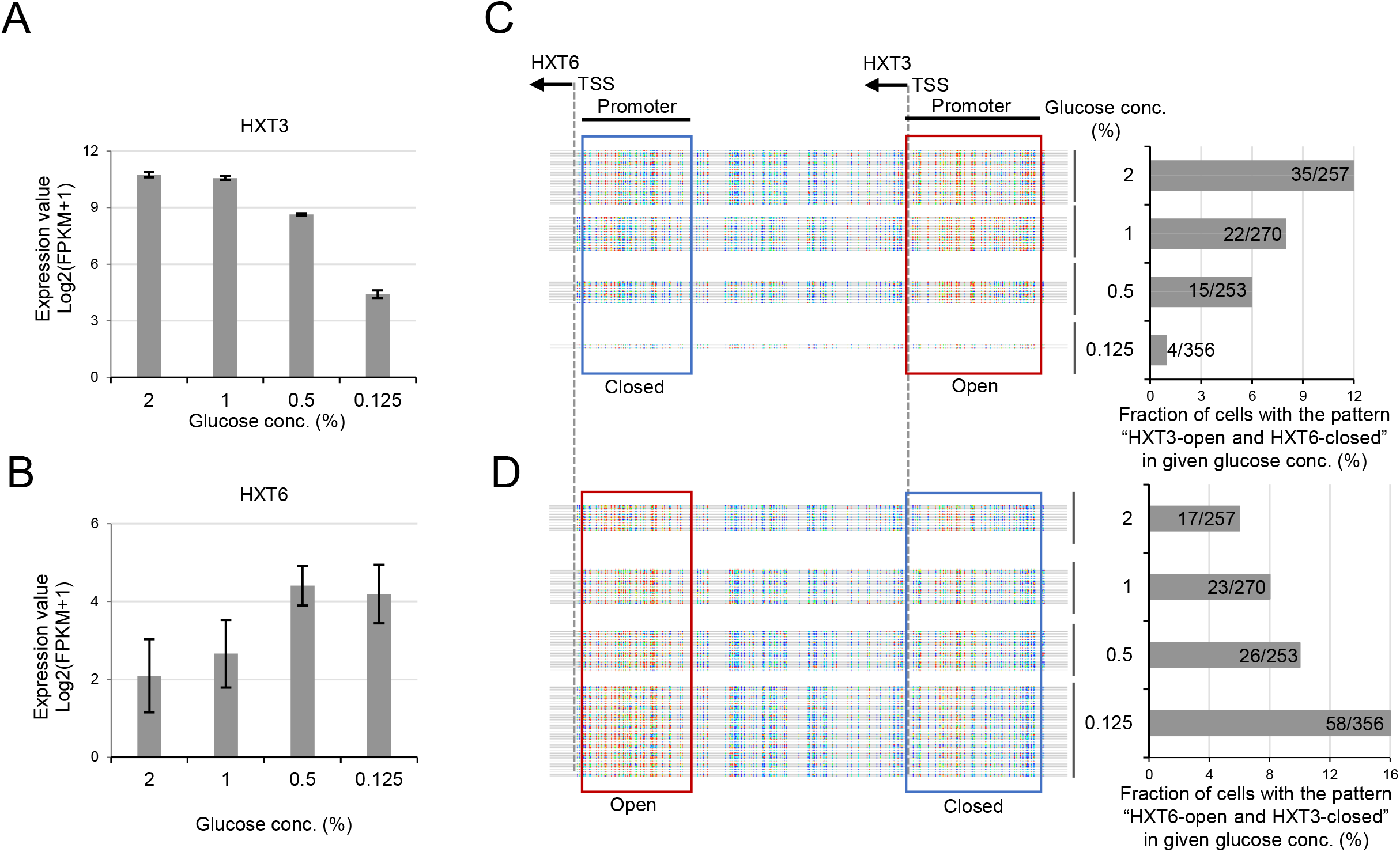
Relationship between chromatin accessibility and co-expression of *HXT3* and *HXT6*. **A, B.** Expression levels of *HXT3* and *HXT6* in response to glucose concentration change. FPKM (Fragment Per Kilobase Million) from bulk-cell RNA-seq data was taken as the expression level. **C, D.** Change of the coupled chromatin statuses of *HXT3* and *HXT6* in response to different glucose concentration (**C**: open-closed; **D**: closed-open). Chromatin accessibility in promoters of *HXT3* and *HXT6* at the single-cell level is shown. Each line represents a molecule (i.e., cell). GpC site is labeled as rainbow-color dot, with methylation score from 0 (blue) to 1.0 (red). A promoter was defined as “open” (highlighted by red box) if the methylation scores of the including GpC sites had a median value greater than 0.5, and “closed” (highlighted by blue box) otherwise. Cells are shown in four groups that corresponded to four glucose concentrations. The cell fractions are also shown on the bar charts.

For each glucose concentration (2%, 1%, 0.5% and 0.125%), we counted MeSMLR-seq molecules to estimate the fractions of cell subpopulations with two opposite coupled chromatin accessibility patterns: “Open-HXT3 and Closed-HXT6” and “Closed-HXT3 and Open-HXT6”. The fraction of cell subpopulation with the coupled pattern “Open-HXT3 and Closed-HXT6” decreased along with the reduction of glucose concentration, whereas “Closed-HXT3 and Open-HXT6” increased (Fig. 10C, D). The changes of two coupled patterns matched the expression dynamics of two genes in response to glucose concentration change (Fig. 10A-D). These proof-of-concept results highlight the promising utility of MeSMLR-seq on studying complex epigenetic changes during transcription reprogramming.

## DISCUSSION

A large number of studies have demonstrated key regulatory roles for nucleosome positioning and chromatin accessibility in eukaryotic gene expression (33–36) as well as DNA repair, recombination and other DNA-dependent processes (37–42). The relationship between nucleosome positioning, chromatin accessibility and gene expression has been studied most extensively (43). However, unlike the well-studied heterogeneity of gene expression based on single-cell analyses, the heterogeneity of nucleosome positioning and chromatin accessibility is poorly studied due to limitations in experimental and sequencing techniques. Previous bulk-cell studies based on the well-developed experimental techniques established the fundamental knowledge base, while their corresponding versions at the single-cell platforms have not yet lead to more details. This is largely due to the sparse sequencing coverage and short read length. MeSMLR-seq provides an alternative way to address this bottleneck: long read length guarantees the full length of genomic region of interest (e.g., whole gene body together with the flanking neighborhood) can be covered by many single reads (that is, single DNA molecules). In the application to haploid organisms, MeSMLR-seq read population represents cell population, so the heterogeneity at the cell level can be investigated. In this study, MeSMLR-seq provides a long-range chromatin status landscape and nucleosome positioning detection at the single-molecule/-cell level. The investigation of coupled chromatin changes and differential nucleosome organization principles in response to nutrition changes underline the unique MeSMLR-seq output on exploring these complex epigenetic events.

However, it should be noted that the molecule-cell link does not hold in diploid or polyploid organisms, as the molecule populations is a mix of allele-specific and cell-specific events. It leads to challenges yet opportunities in the further development of new experimental (e.g., single-cell barcoding) and statistical (e.g., data deconvolution) approaches. Once cell subpopulations can be reconstructed from a molecule population, we could distinguish the allele-specific epigenome precisely from different cell subpopulations and achieve more accurate investigation of how epigenetics events behave differently at alleles. Regardless of the wide interest on the cell-level study, the characterization of nucleosome positioning and chromatin status at single DNA molecules by MeSMLR-seq will also bring very unique and informative data to reveal the dynamic nucleosome positioning mechanism, such as assembly, disassembly, and sliding.

Besides the single-molecule information, the long length of MeSMLR-seq reads, which allows correlation analysis of exogenous and endogenous methylation statuses over different positions, could be informative for some research topics: 1) correlation of exogenous 5mC events has shown the nucleosome positioning pattern in this study (Fig. 2B), and thus DNA loops or other larger spatial chromatin domain that affects exogenous methylation could be also identified, which would require specific library preparation to generate even longer ONT reads; 2) As endogenous 5mC can be also detected, MeSMLR-seq can be applied to other higher organisms (e.g., human) to study how methylation status at different genomic region coordinates, but it could also provide direct evidence to address the controversial topics about how methylation status and nucleosome positioning and chromatin openness correlates.

On a technical view, there is relatively few application of ONT data at epigenetics research, as the corresponding experimental approaches or bioinformatics methods are rarely developed, although numerous applications of ONT data have been published rapidly with improved data quality and cost efficiency. In addition to the previously reported studies of identifying methylation and three-dimensional spatial organization of chromatin (44), MeSMLR-seq contributes a new technique in the toolkit of single-molecule long-read sequencing to obtain the first-hand details of epigenetics at single DNA molecules. More other innovative studies with single-molecule long-read sequencing should be explored and expected to advance our studies to discover novel and complex biological insights.

## MATERIALS AND METHODS

### Yeast strain and growth

*Saccharomyces cerevisiae* BY4741 strain was used in this study. Yeast cells were separately grown at 30°C in the media including 1% yeast extract, 2% peptone and different carbon sources. Yeast cells were collected in the mid-log phase (OD_600_ of 0.3-0.6) and subjected to MeSMLR-seq, bulk-cell RNA-seq and single-cell RNA-seq experiments (SI Appendix, section 4, section 5 and Table S4).

### MeSMLR-seq experiment

Preparation and methylation of yeast spheroplasts were performed as previously described (16) (Fig. 1 and SI Appendix, Fig. S1). Briefly, yeast cells were treated with Zymolyase (amsbio, final conc. = 0.25 mg/mL) in 1 M sorbitol and 50 mM Tris (pH7.4) and 10 mM β-mercaptoethanol. Spheroplasts were washed using 1 M sorbitol twice before methyltransferase treatment. GpC-specific methyltransferase M.CviPI (NEB) supplemented with 160 μM SAM S-adenosylmethionine was used to methylate spheroplasts at 37°C for 45 min. Genomic DNA was extracted using PCI (Phenol:chloroform:isoamyl alcohol, 25:24:1) and purified by Genomic DNA Clean & ConcentratorTM-10 Kit (Zymo Research).

We denote the above mentioned genomic DNA that undergoes *in vivo* spheroplast methylation as target sample of MeSMLR-seq. Native genomic DNA extracted from yeast without M.CviPI treatment was used as negative control (all cytosines at GpC sites are unmethylated). There is no endogenous 5mC in yeast genome as reported in previous study (20). Genomic DNA treated by M.CviPI (without spheroplast methylation) was used as positive control (all cytosines at GpC sites are 5mCs).

The efficiency of M.CviPI methylation was evaluated using bisulfite sequencing as previously described (16). Firstly, bisulfite conversion was performed using EZ DNA Methylation Lighting Kit (Zymo Research). Secondly, PCR amplification targeted to specific genomic regions was performed by ZymoTaq PreMix (Zymo Research). *CHA1* gene region (ChrIII:15713-16074), *CYS3* gene region (ChrI:130966-131117), *GAL10* gene region (ChrII:278464-278738) and *PHO5* gene region (ChrII:430248-430388) were amplified for evaluating the methylation efficiency of positive control. The *PHO5* gene region (ChrII:430843-431498), which was shown in the Figure 1 of the previous study (16), was used to estimate the efficiency of spheroplast methylation (i.e., target sample of MeSMLR-seq). Thirdly, TA cloning was performed by TOPOTM TA Cloning Kit (Life Technologies). Single colonies were picked up and plasmids were extracted by QIAprep Spin Miniprep Kit (QIAGEN). Finally, plasmids were sequenced by Sanger sequencing. For positive control, we estimated the efficiency of methylation as the percentage of 5mC over all GpC sites (totally 53 GpC sites for four target gene regions). Three single colonies were sequenced per gene region; and the methylation efficiency of positive control was ((53×3)-1)/(53×3) = 99.37%. For target sample of MeSMLR-seq, we considered it as successfully-methylated if the single colony included at least one 5mC. In total, 13 colonies were sequenced; and the methylation success rate of target sample was up to 100% (13/13).

Native genomic DNA (negative control), methylated genomic DNA (positive control) and extracted genomic DNA after spheroplast methylation (target sample) were directly submitted to ONT sequencing. In brief, the genomic DNA was fragmented (size = 8 kb) using Megaruptor. Sequencing library was prepared using the 1D Ligation Sequencing Kit (SQK-LSK108). ONT sequencing was performed using GridION platform with R9.4.1 flow cells.

### MeSMLR-seq data preprocessing

The base-called ONT sequencing data were aligned to sacCer3 reference genome using BWA software (version 0.7.17-r1188) (45) with the “mem” mode and the “-x ont2d” parameter. Nanopolish (version 0.8.5) (18) with the “eventalign” mode and the “--scale-events” parameter was used to generate the alignments between event levels and 6-mers for each sequencing molecule, which were utilized for the following GpC-specific 5mC detection.

Since we used ONT 1D sequencing strategy in this study, a DNA molecule from yeast cell might be sequenced twice (i.e., forward and reserve strands). Thus, to achieve the “one-to-one” link between ONT sequencing molecule and haploid yeast cell, we classified all molecules into two groups based on their aligned genomic strands: forward and reverse.

The MeSMLR-seq data was summarized in SI Appendix, Table S1.

### GpC-specific 5mC detection at the single-molecule level and single-base resolution by MeSMLR-seq

For every unique 6-mer (4^6^=4096 in total), we modeled the event level for unmethylated cytosine by a Gaussian distribution, and the event level for methylated cytosine 5mC at GpC site by a Gaussian mixture distribution considering the fact the efficiency of exogenous methylation was not always 100% (99.37% in our experiment) (SI Appendix, Fig. S2A, right panel). Based on the native and positive control data, the corresponding distribution parameters were estimated by the sample mean and standard variation, and by the EM algorithm (for the Gaussian mixture model), respectively. The area of the overlapped region under the two probability density functions was calculated. The discrimination of a given 6-mer was defined as (1 - the area of overlap).

Given a GpC site on the reference genome and a sequencing molecule from target sample, we listed all the 6-mers that covered the cytosine at GpC dinucleotide (SI Appendix, Fig. S2A, left panel). The 6-mer with >1 GpC site or >10 aligned event levels from the molecule was excluded for 5mC detection. Among the remaining 6-mers, the one with the maximal discrimination was chosen for the calculation of methylation score.

Denote *k* as the selected 6-mer, and *e* as the event level that is aligned to *k*. Let *f_P_*(*e; k*) and *f_N_*(*e; k*) be the values of probability density functions for 5mC and unmethylated cytosine, respectively. The event level *e* was filtered out if one of log*f_P_*(*e; k*) or log*f_P_*(*e; k*) was <-10; otherwise, the methylation score of the GpC site is calculated as

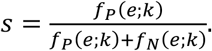

The score *s* is essentially the posterior of methylation given a non-informative prior. If multiple event levels were aligned to *k*, then *f_P_*(*e; k*) and *f_P_*(*e; k*) were replaced by the product of the multiple likelihood.

To evaluate the performance of 5mC detection, we plot receiver operating characteristic (ROC) curve (Fig. 2A). In detail, the negative control and positive control data were randomly split into two sets with equal size, respectively. One of them was used for training, and the other for test.

### Nucleosome positioning detection at the single-molecule level by MeSMLR-seq

We developed a bioinformatics method, named **NP-SMLR** (**N**ucleosome **P**ositioning detection by **S**ingle-**M**olecule **L**ong-**R**ead sequencing), to detect and phase nucleosomes at the single-molecule level (Fig. 2C).

Let *X*_1_*X*_2_ … *X_l_* be a molecule, where *X_i_* is the *i*-th base. Denote *s_i_* as the methylation score of *X_i_*, if *X_i_* is the cytosine of the GpC dinucleotide. Suppose that the event levels of all GpC sites are independent. Nucleosome positioning detection refers to finding a path ***π*** = *π*_1_*π*_2_ … *π_l_* that maximizes the likelihood of signals:

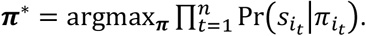

*π_i_* takes the value from {*L, N*_1_, *N*_2_, …, *N*_147_}. *L* represents the linker region; *N_m_* represents the *m*-th base within a nucleosome; *i*_1_, *i*_2_, …, *i_n_* are the positions of cytosines that belong to GpC dinucleotides. The elements of path *π* are restricted that: 1) *N_m_* is followed by *N_m+t_* (1 ≤ *m* ≤ 146); 2) *N*_147_ is followed by *L*; and 3) *L* is followed by *L* or *N*_1_.

Based on the methylation scores of all GpC sites from all molecules in negative and positive control training data, we can fit two density curves using the “density” command in R (version 3.3.0), respectively. The two density functions are denoted as *q_N_*(·) and *q_P_*(·), respectively (SI Appendix, Fig. S2B). A dummy methylation score *s_i_* = −1 is added for *X_i_* if it is not a cytosine of GpC dinucleotide. Define

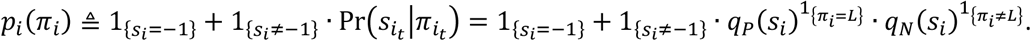

Let *a_π_i_,π_i+1__* be the compatibility indicator of two adjacent states such that

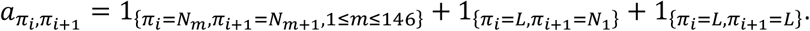

The objection function can therefore be expressed as

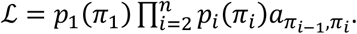

Define

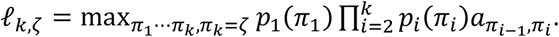

Then the maximum of objection function can be obtained by iteration:

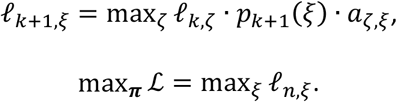

Accordingly, ***π***\* can be obtained through dynamic programming (Fig. 2C). We start by building an *l* × 148 matrix *V*. Line *i* corresponds to *X_i_*, the *i*-th base of the molecule. Column 1 corresponds to the linker; and the other columns (from Column 2 to Column 148) correspond to *N*_1_, *N*_2_, …, *N*_147_, separately. Initialize *V*[1,1] = *p*_1_(*L*), and *V*[1, *j*] = *p*_1_(*N*_*j*−1_), 2 ≤ *j* ≤ 148. Elements in Line *i*(2 ≤ *i* ≤ *n*) are then calculated iteratively. For Column 1, the element *V*[*i*, 1] is set as max{*V*[*i* − 1,1], *V*[*i* − 1,147]}*q_P_*(*s_i_*) if *X_i_* is cytosine of GpC, or max{*V*[*i* − 1,1], *V*[*i* − 1,147]} otherwise. For Column *j* (2 ≤ *j* ≤ 148), *V*[*i,j*] is set as *V*[*i* − 1, *j* − 1]*q_N_*(*s_i_*), or *V*[*i* − 1, *j* − 1] otherwise. When updating an element, we record the position of the previous element that leads to the maximal value, and store the position as a pointer. After updating all elements, the maximal element in the last line is found (elements that equal to one are not considered), and the nucleosome positioning detection is completed through the backtracking of pointers. All calculations are performed in log scale to avoid rounding error.

We evaluated the accuracy of nucleosome positioning detection (NP-SMLR) through simulation tests under different nucleosome coverage and GpC frequency (Fig. 2D). In detail, DNA sequence (3-kb length) was simulated with randomly assigned GpC sites at given frequency. Lengths of linkers between nucleosomes were sampled independently and sequentially. At each time, the linker length was sampled from the normal distribution 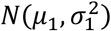 with probability *τ*, and 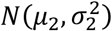 with probability 1 − *τ*, corresponding to regular nucleosome array and open region with specific biological functions, respectively. We set *μ*_2_ > *μ*_1_ and 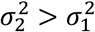. Nucleosomes were then placed on the DNA sequence, with their distance being set as the above simulated linker length. Methylation scores for GpC sites occupied by nucleosomes were generated based on the score distribution of negative control data, whose density function was *q_N_* (·). For GpC sites within linkers, *q_P_*(·) was used instead. NP-SMLR was applied on the simulated sequence. Denote 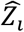 and *Z_i_* as the predicted and real indicators of whether the *i*-th base locates in nucleosome or not, respectively. The accuracy was defined as

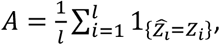

where *l* is the length of the simulated DNA sequence. In simulation tests, we set *μ*_1_ = 15, *σ*_1_ = 5, *σ*_2_ = 10, *τ* = 0.1. We set *μ*_2_ as 15, 50, 100, 200, 300, 400, 500, 600, respectively to achieve different nucleosome coverage (defined as the proportion of bases covered by nucleosomes). For each parameter setting, the above simulation was carried out for 1,000 times.

### Bulk-cell level nucleosome occupancy analyses based on MeSMLR-seq data

The genomic coordinates of all nucleosomes predicted by NP-SMLR at the single-molecule level were pooled and subjected to iNPS software (version 1.2.2) (46) with default parameters to generate bulk-cell level nucleosome occupancy profile and to call nucleosome peaks.

The nucleosome occupancy profiles were used to generate Fig. 3A, B; Fig. 4D, E (upper panel); Fig. 7A, B; and SI Appendix, Fig. S6D, E. The nucleosome peaks called by iNPS were used for the comparison with MNase-seq (Fig. 3C).

### Measurement of nucleosome positioning heterogeneity

The heterogeneity of nucleosome positioning was measured by the variation of the +1 nucleosome positioning relative to TSS across different cells (Fig. 4B and SI Appendix, Fig. S3A). For each molecule/cell, we first defined the nucleosome whose center was located in the downstream of TSS and closest to TSS as the +1 nucleosome. Next, we sorted the distances between +1 nucleosomes and TSS, and removed the upper 10% values for robustness. The standard variance of the remaining values was used to represent the heterogeneity of nucleosome positioning for each gene.

### Measurement of nucleosome spacing uniformity

The uniformity of nucleosome spacing was measured by the variation of the distance between adjacent nucleosomes (i.e., the length of linker region) (Fig. 4C and SI Appendix, Fig. S3B). For each gene, the molecules that fully covered the region (from upstream 500 bp to downstream 100 bp of TSS) were chosen. For each molecule, we calculated the lengths of all linker regions that were located in the region “-500, +100”. Then, we calculated the absolute deviation of linker length pair-wisely. The sum of the deviation values was divided by the number of linker pairs. The obtained value, which described the variation of nucleosome distance, was namely the nucleosome spacing uniformity.

### Chromatin accessibility mapping at the single-molecule level based on MeSMLR-seq data

Based on the methylation scores of all GpC sites per molecule, we detected accessible chromatin regions along the molecule. Given a single molecule *X*_1_*X*_2_ … *X_l_*, where *X_i_* is the *i*-th base, we defined the interval from *X_i_* to *X_j_* as an accessible region if: 1) *X_i_* and *X_j_* were adjacent GpC sites; 2) the corresponding methylation scores *s_i_* and *s_j_* were >0.5; and 3) the distance between *X_i_* and *X_j_* was <100 bp. The continuous accessible regions were merged. Given an accessible region, the chromatin accessibility score was defined as the median methylation score among all GpC sites within this region.

In this study, we only considered the accessible regions with the length ≥100 bp for each molecule. Genome-wide chromatin accessibility profile was generated through merging accessible regions of all molecules. The chromatin accessibility profile was used to generate the Fig. 5A, B; Fig. 6 (upper panel); Fig. 7C; Fig. 8A; SI Appendix, Fig. S5A, B, and SI Appendix, Fig. S6B, C.

### Chromatin accessibility peak calling at the bulk-molecule/-cell level based on MeSMLR-seq data

We defined significantly-accessible genomic regions as described in the previous study (30). Let *G_i_* be the *i*-th base of the genome. Denote 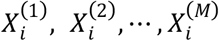 as the bases from *M* sequencing molecules that covered *G_i_*, and 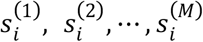 as the corresponding methylation scores if *G_i_* is a GpC site. Define 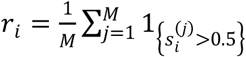, which is the ratio of methylated bases (methylation score >0.5), and denote 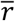 as the average of ratios of all GpC sites. We defined the interval between *G_i_* and *G_j_* as a significantly-accessible region if: 1) *G_i_* and *G_j_* were adjacent GpC sites; 2) 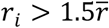, and 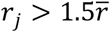; and 3) the distance between *G_i_* to *G_j_* was <100 bp. The continuous accessible regions were merged to generate a longer accessible genomic region (referred as “chromatin accessibility peak”).

In this study, we only considered the peaks with the length ≥100 bp. For sequencing molecules aligned to forward and reverse genomic strands, we defined chromatin accessibility peaks, separately. The overlapped peaks between the forward and reverse strands were used for the comparison with two existing methods (i.e., ATAC-seq and DNase-seq) (Fig. 5C).

### Definition of gene promoter region and measurement of gene accessibility

To quantitatively measure the accessibility of genes, we first defined the promoter region for each gene. Briefly, chromatin accessibility peaks (including both forward and reverse strands) were called using MeSMLR-seq data for each biological sample. For each biological sample, the overlapped peaks between forward and reverse strands for MeSMLR-seq were merged together. Next, we combined the merged peaks of MeSMLR-seq from all biological samples and the overlapped peaks between two biological replicates of DNase-seq. For each gene, 1) if there was only one peak that was located within the upstream 500 bp and downstream 100 bp of TSS (named “-500, +100” region), the peak was defined as the promoter region; or 2) if there were multiple peaks that were located in the “-500, +100” region, the peak that had the longest overlap was defined as the promoter region; or 3) if there was no peak locating in the region “-500, +100”, the region “-500, +100” was defined as the promoter region.

At the single-molecule level, the accessibility score of a gene was calculated as the median methylation score among all GpC sites within the promoter region. For all molecules covering the promoter of a given gene, we categorized them into two chromatin statuses: “open” if the accessibility score was >0.5; “closed” otherwise. The defined promoter region and the corresponding accessibility score were used to generate the Fig. 5E; Fig. 6 (upper panel); Fig. 8D; Fig. 9; Fig. 10C, D; and SI Appendix, Fig. S6A.

### Analyses of dynamic gene expression and chromatin accessibility among three carbon sources

Differentially-expressed genes were identified using Cuffdiff (*q*-value <0.01) between glucose (Glu) and other two carbon sources, galactose (Gal) and raffinose (Raf). Overall, there were 700 up-regulated and 682 down-regulated genes in Gal (Glu *vs*. Gal); 605 up-regulated and 727 down-regulated genes in Raf (Glu vs. Raf). These differentially-expressed genes were used to generate the Fig. 8B-D. Gene enrichment analyses in Fig. 8C was performed using DAVID (version 6.8) (50).

For the differential chromatin accessibility analyses, we first calculated the bulk-cell-level chromatin accessibility as the ratio of those with “open” status among the molecules that fully covered the gene promoter. For each gene, the differential chromatin accessibility score was computed as the difference of bulk-cell-level chromatin accessibility between two carbon sources (Glu minus Gal for “Glu vs. Gal”; Glu minus Raf for “Glu vs. Raf”).

## ACKNOWLEDGEMENTS

We would like to thank Dr. Charles Brenner (Department of Biochemistry, University of Iowa) for providing yeast BY4741 strain. We are grateful to Drs. David Stoltz and Massimo Attanasio (Department of Internal Medicine, University of Iowa) for experimental support.

This work was supported by an institutional fund of Department of Internal Medicine, University of Iowa and an institutional fund of Department of Biomedical Informatics, The Ohio State University (to K.F.A., Y.W., A.W., and Z.L.), the National Institutes of Health (R01HG008759 to K.F.A., Y.W., A.W., and Z.L.), and the Multidisciplinary Lung Research Career Development Program (T32HL007638 to A.T.).

## Supplementary Information

### Section 1: Analyses of ATAC-seq, DNase-seq, MNase-seq, ChlP-seq, ChlP-exo and ChEC-seq data

The information (including yeast strain, growth condition, GEO accession number, data format and reference) of public sequencing data used in this study was summarized in SI Appendix, Table S6.

Quality control of raw sequencing data (FASTQ format) was performed using FastQC and cutadapt; and alignment was performed using Bowtie2 software (version 2.2.5) (1) with default parameters.

For ATAC-seq (2) and ChIP-seq (Pol2) (3) data, MACS2 software (version 2.2.1) (4) with default parameters was used to call significantly-enriched peaks (*q*-value <0.05).

For MNase-seq data (5), iNPS with default parameters was used for nucleosome calling.

For DNase-seq data (6), F-Seq software (version 1.85) (7) with default parameters was used to call significantly-enriched peaks (peak length ≥100 bp).

For ChIP-exo (Abf1, Cbf1, Mcm1, Rap1 and Reb1) data, the called peak files were directly downloaded from the original study (8).

For ChEC-seq (Med8 and Med17) data (9), chec-seq script (https://github.com/zentnerlab/chec-seq) was used to call significantly-enriched peaks (signal-noise ratio ≥10 and peak length ≥100 bp).

### Section 2: Correlation and overlapping analyses between MeSMLR-seq and MNase-seq

For correlation analysis of the bulk-cell level nucleosome occupancy results, we used iNPS to generate nucleosome occupancy profiles (BigWig format) for MNase-seq and MeSMLR-seq, respectively. Pearson correlation coefficient of nucleosome occupancy profiles (across whole genome and bin size as 10 bp) was calculated between two methods (Fig. 3A).

For overlapping analysis of nucleosomes, we only considered the two nucleosome peaks (from MeSMLR-seq and MNase-seq, respectively) as overlapped if ≥50% region of one peak was covered by another peak (Fig. 3C).

### Section 3: Correlation and overlapping analyses among MeSMLR-seq, ATAC-seq and DNase-seq

For correlation analysis of the bulk-cell level chromatin accessibility results, we generated genome-wide chromatin accessibility profiles (BigWig format) for three methods, separately. Pearson correlation coefficient of chromatin accessibility profiles (across the whole genome and bin size of 10 bp) were calculated among three methods (Fig. 5A).

For MeSMLR-seq data, we separately called significantly-enriched peaks for molecules aligned to forward and reverse strands. Only the overlapped peaks between the forward and reverse strands for MeSMLR-seq data, and the overlapped peaks between two biological replicates for ATAC-seq and DNase-seq were used for overlapping analysis (Fig. 5C).

### Section 4: Single-cell RNA-seq experiment and data analysis

Yeast cells growing in YPD (1% yeast extract, 2% peptone and 2% glucose) medium were collected and spheroplasts were prepared as described above. Cell viability was measured using Trypan blue exclusion method and cell number was counted by hemocytometer. Of note, considering the fragility of spheroplasts, we modified the loading strategy of buffer before running the 10X ChromiumTM Controler (10X Genomics). Firstly, Single Cell Master Mix (10X Single Cell 3’ Reagent Kit v2) was prepared and added into Single Cell A Chip. Next, instead of nuclease-free water, sorbitol was added (final conc. = 1 M) and mixed well. Finally, spheroplasts suspended in 1 M sorbitol were added. In total, 318 million read pairs (2 × 150 bp) were generated by Illumina HiSeq 4000 platform.

The quality of single-cell RNA-seq (scRNA-seq) data was evaluated by FastQC software. Cellranger software (version 2.1.1) with default parameters was used to process scRNA-seq data and generate gene-cell matrix. For quality control of scRNA-seq data, we excluded the cells with >10,000 UMI (unique molecular identifier) counts as they were potentially from artificial cell or cell duplets (10). After quality control, 2,812 single cells with 4,335 UMI counts (median value) per cell and 103,002 read pairs (median value) per cell were used in the following analyses. The number of expressed genes (≥1 UMI) per cell was 1,572 (median value). DESeq2 package (11) was used to normalize scRNA-seq UMI count data for 2,812 cells.

### Section 5: Bulk-cell RNA-seq experiment and data analysis

Total RNA was extracted using Quick-RNA Fungal/Bacterial Miniprep Kit (Zymo Research). Sequencing library was prepared using TruSeq Stranded mRNA Library Prep Kit and 10 million read pairs (2 × 150 bp) on average per sample were generated using Illumina HiSeq 4000 platform. Three biological replicates per biological condition were performed.

The quality of bulk-cell RNA-seq data was evaluated by FastQC software (version 0.11.3, https://www.bioinformatics.babraham.ac.uk/projects/fastqc/) and sequencing adaptors were trimmed by Cutadapt software (version 1.8.1) (12). Processed reads were aligned to reference genome (version UCSC sacCer3) by Hisat2 software (version 2.0.0-beta) (13) with default parameters. Cufflinks (version 2.2.1) (14) with default settings were separately used for quantifying gene expression, normalizing gene expression and analyzing differential gene expression. The cutoff of statistical significance of differential gene expression was *q*-value < 0.01.

The bulk-cell RNA-seq data was summarized in the SI Appendix, Table S5.

**Fig. S1.**
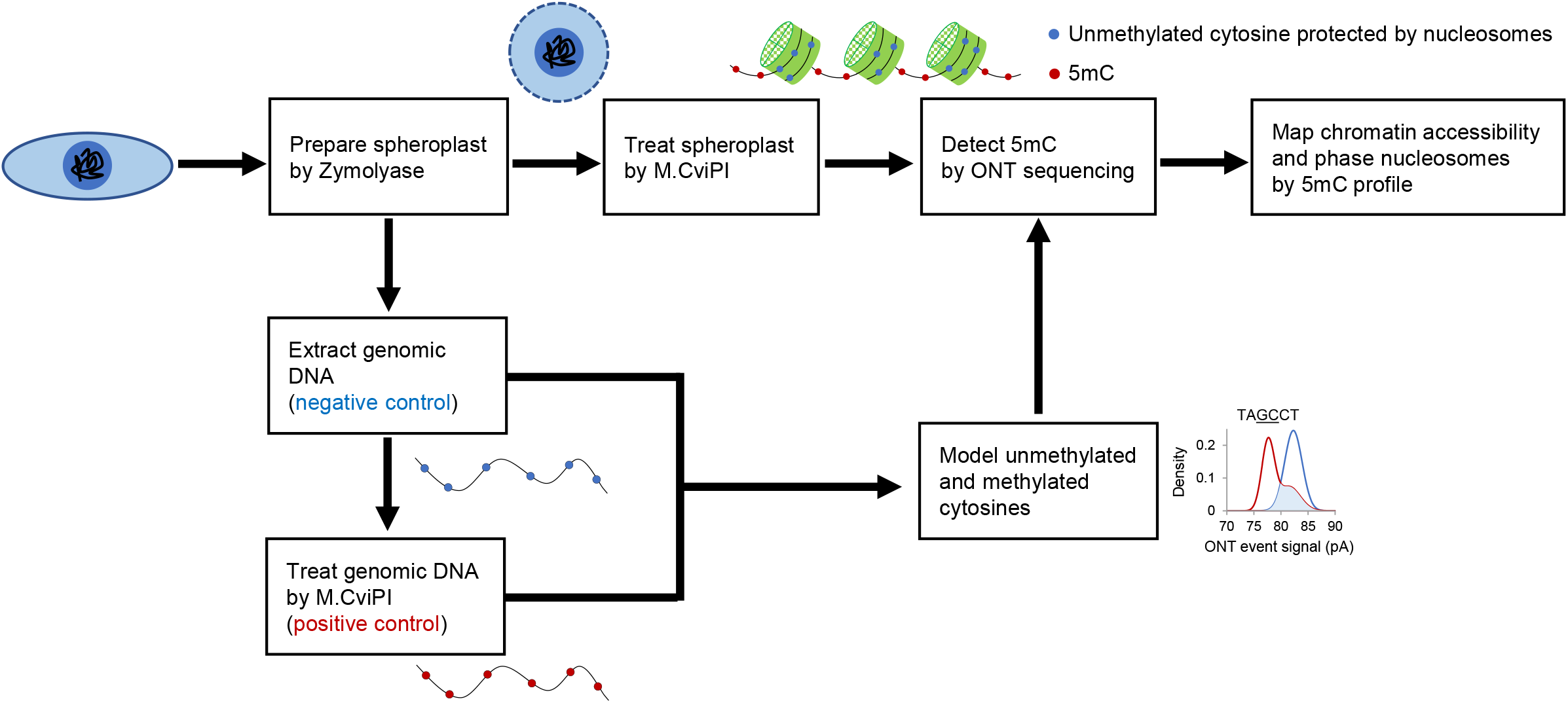
Flowchart of MeSMLR-seq

**Fig. S2.**
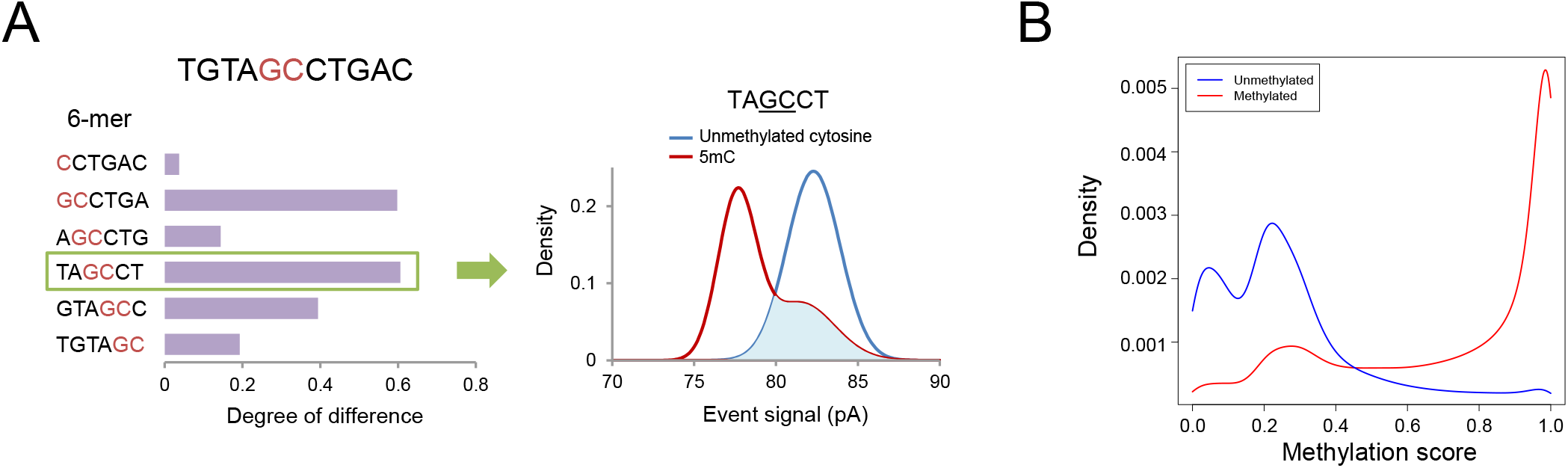
5mC methylation calling at GpC sites and distribution of methylation scores. **A.** An example showing the difference on event level distribution of a 6-mer with unmethylated cytosine or 5mC at GpC site (right panel). Among all 6-mers covering a GpC site, the one with the largest degree of difference was chosen for methylation detection (left panel). **B.** The probability distribution of methylation scores for negative and positive control data. The figure was drawn based on the data that were used for 5mC detection test.

**Fig. S3.**
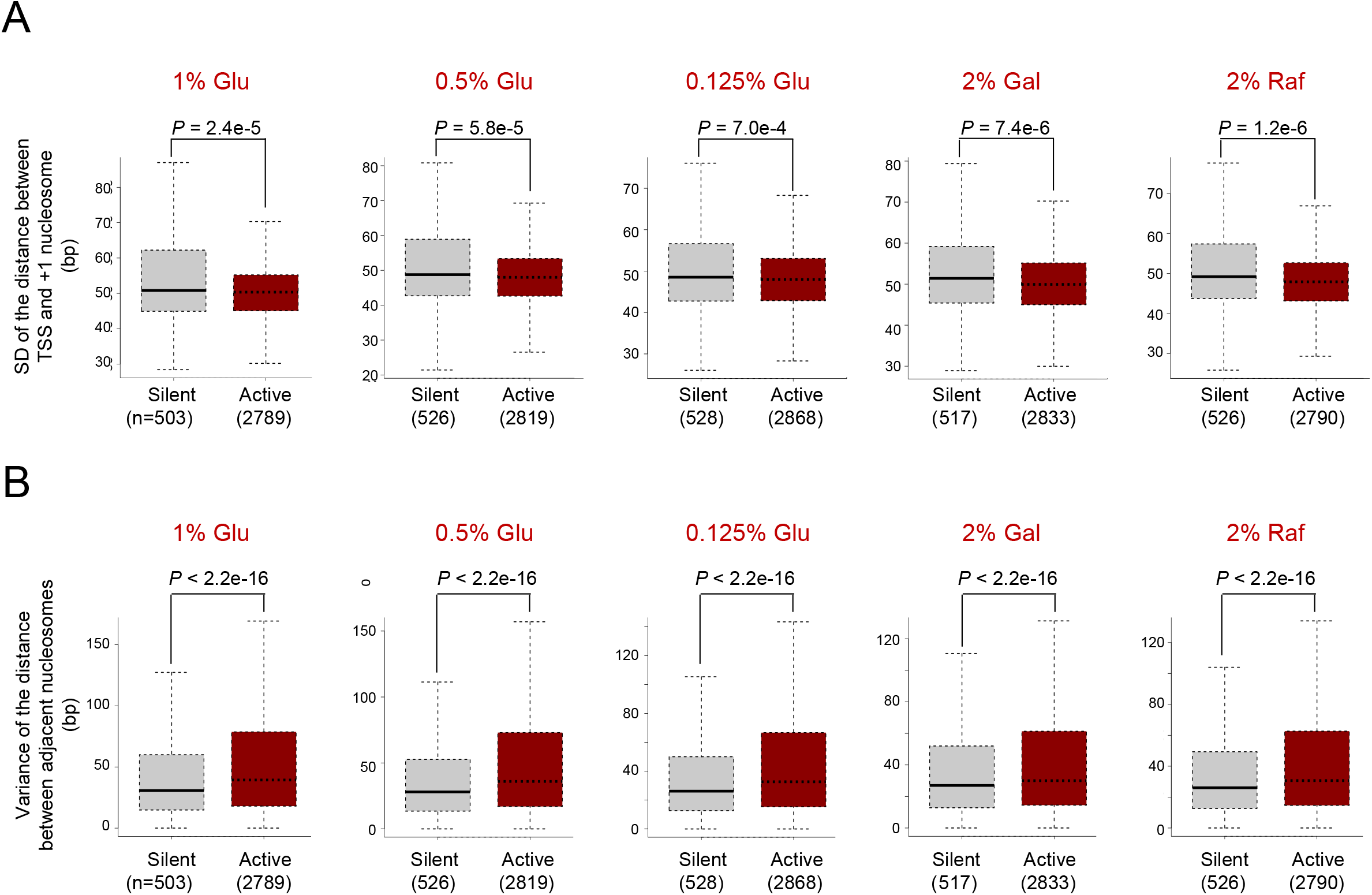
Heterogeneity of nucleosome positioning and uniformity of nucleosome spacing. **A.** Heterogeneity of nucleosome positioning for five growth conditions. The heterogeneity of nucleosome positioning was measured by the standard deviation of the distances between +1 nucleosome and TSS. SD, standard deviation. The p-value was calculated by Wilcoxon rank sum test. **B.** Uniformity of nucleosome spacing for five growth conditions. The p-value was calculated by Wilcoxon rank sum test.

**Fig. S4.**
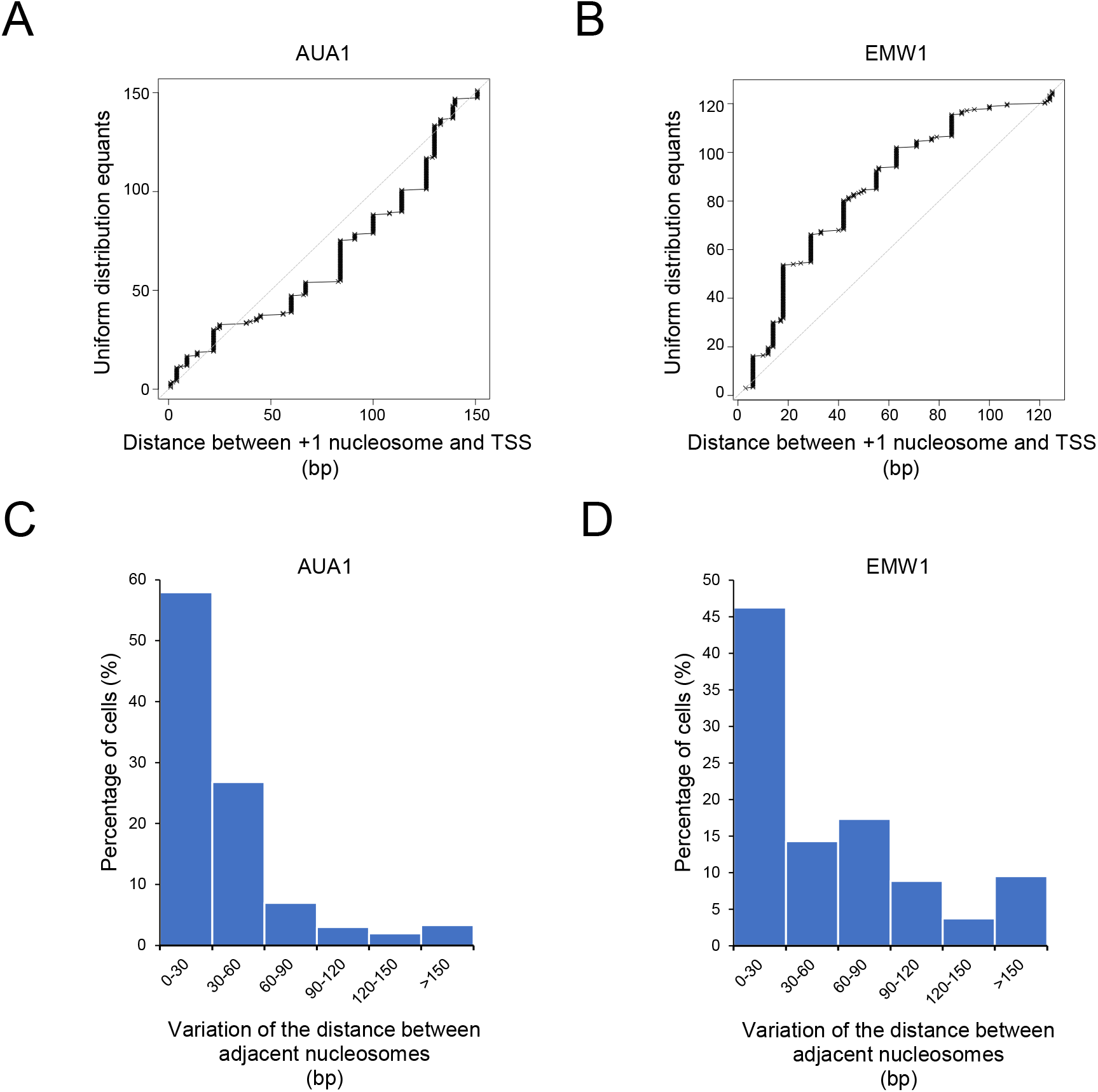
Differential nucleosome organization between silent (*AUA1*) and active (*EMW1*) genes. **A, B.** Q-Q plot illustration of the heterogeneity of nucleosome positioning. Each cross mark represents a molecule/cell. The x-axis is the distance between +1 nucleosome and TSS. The y-axis is the equant under the assumption that all distance values are evenly distributed. **C, D.** Uniformity of nucleosome spacing. Smaller variation (x-axis) indicates that nucleosomes are more likely to be uniformly spaced.

**Fig. S5.**
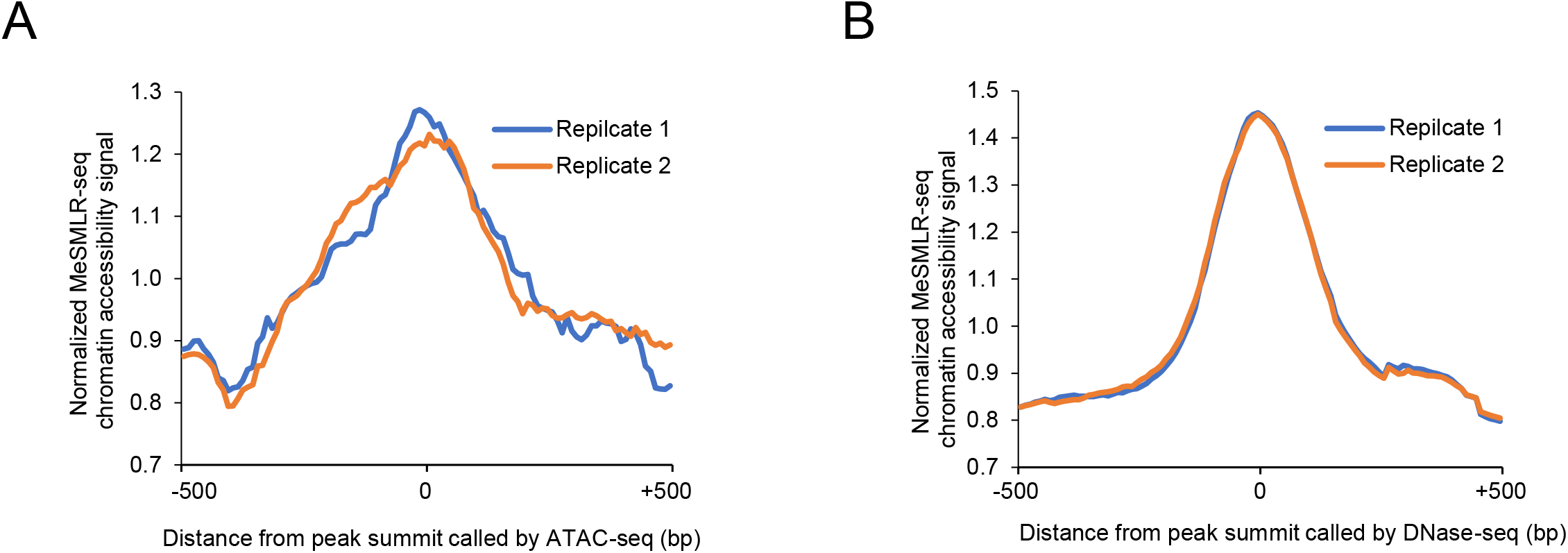
MeSMLR-seq chromatin accessibility signal distribution surrounding the peak summits called by ATAC-seq (A) or DNase-seq (B)

**Fig. S6.**
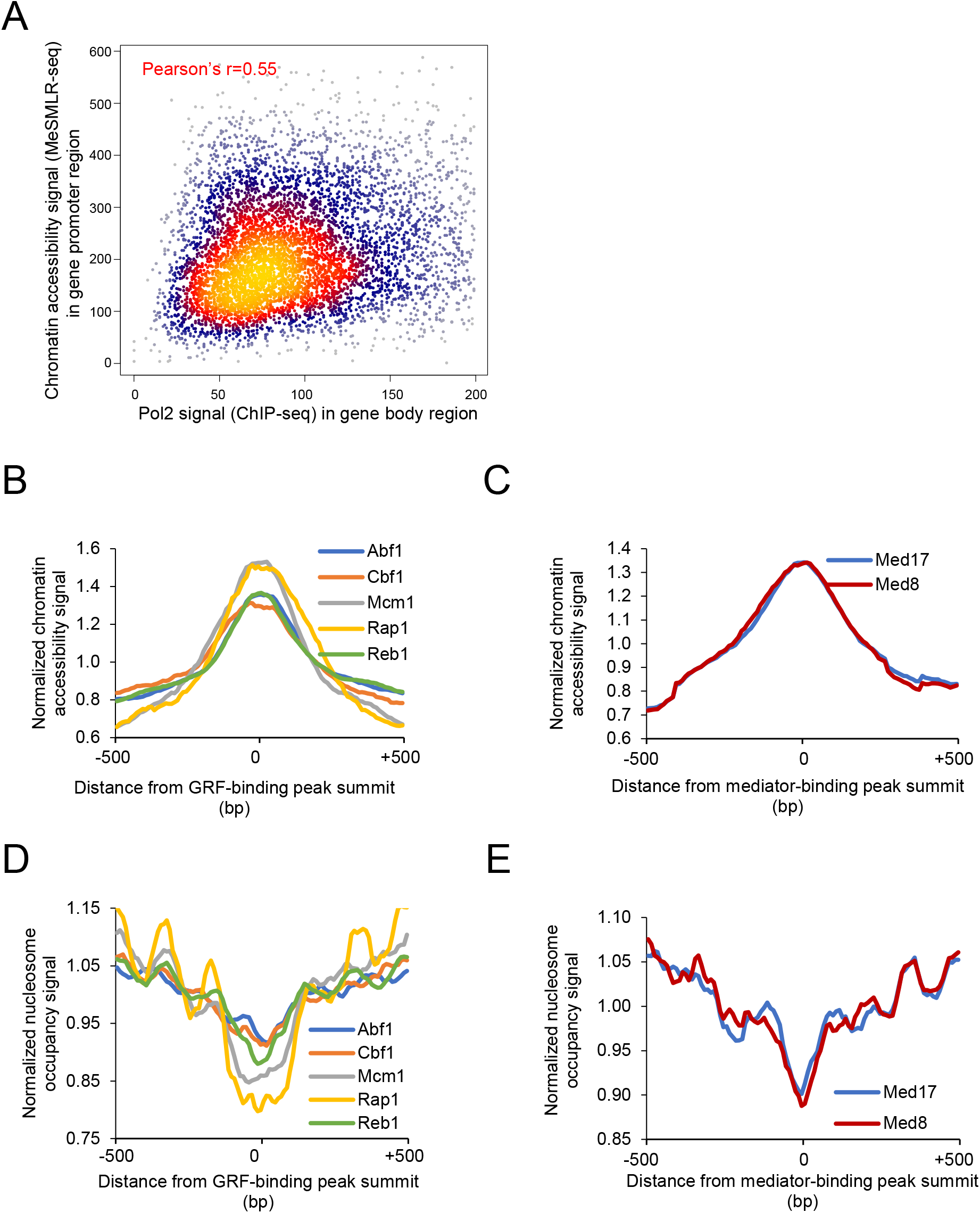
Chromatin accessibility and nucleosome occupancy profiles at the binding sites of transcription-related factors. **A,** Correlation between chromatin accessibility in promoter and Pol2 binding signal in gene body. Each point represents one gene. **B, D.** Chromatin accessibility (**B**) and nucleosome occupancy (**D**) profiles at the binding sites of five general regulatory factors. **C, E.** Chromatin accessibility (**C**) and nucleosome occupancy (**E**) profiles at the binding sites of two mediators.

**Table S1.**
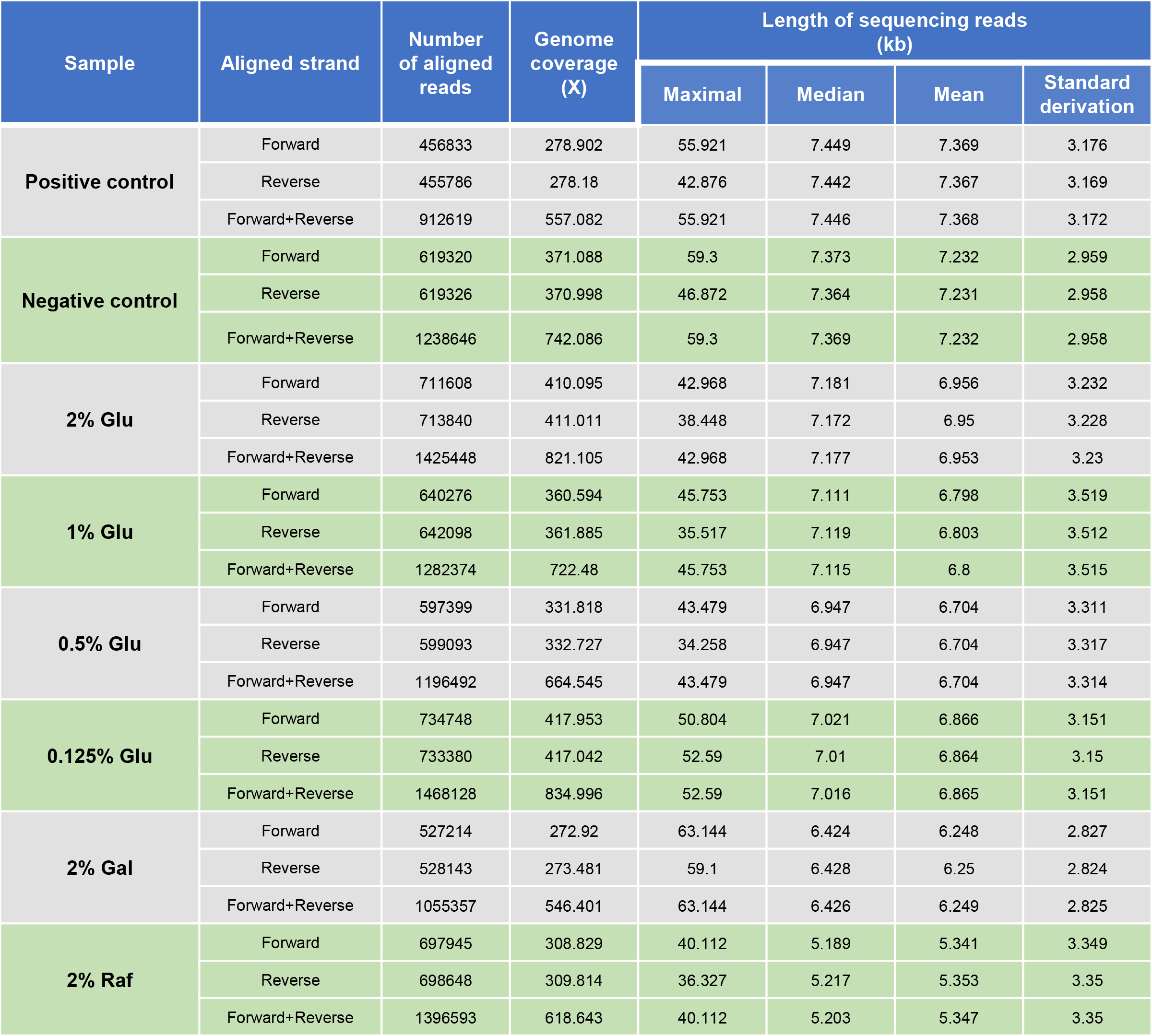
Statistics of MeSMLR-seq data

**Table S2.** Number of nucleosomes phased by single molecules of MeSMLR-seq

| Sample | Aligned strand | Number of genes covered by single molecules |  |  |  |
| --- | --- | --- | --- | --- | --- |
|  |  | Maximal | Median | Mean | Standard derivation |
| 2% Glu | Forward | 244 | 37 | 35 | 18 |
|  | Reverse | 226 | 37 | 36 | 18 |
|  | Forward+Reverse | 244 | 37 | 35 | 18 |
| 1% Glu | Forward | 271 | 36 | 34 | 19 |
|  | Reverse | 207 | 36 | 34 | 19 |
|  | Forward+Reverse | 271 | 36 | 34 | 19 |
| 0.5% Glu | Forward | 256 | 36 | 34 | 19 |
|  | Reverse | 177 | 36 | 34 | 19 |
|  | Forward+Reverse | 256 | 36 | 34 | 19 |
| 0.125% Glu | Forward | 294 | 37 | 36 | 18 |
|  | Reverse | 258 | 37 | 36 | 18 |
|  | Forward+Reverse | 294 | 37 | 36 | 18 |
| 2% Gal | Forward | 306 | 32 | 31 | 16 |
|  | Reverse | 356 | 32 | 31 | 16 |
|  | Forward+Reverse | 356 | 32 | 31 | 16 |
| 2% Raf | Forward | 208 | 26 | 27 | 18 |
|  | Reverse | 199 | 26 | 28 | 18 |
|  | Forward+Reverse | 208 | 26 | 27 | 18 |

**Table S3.**
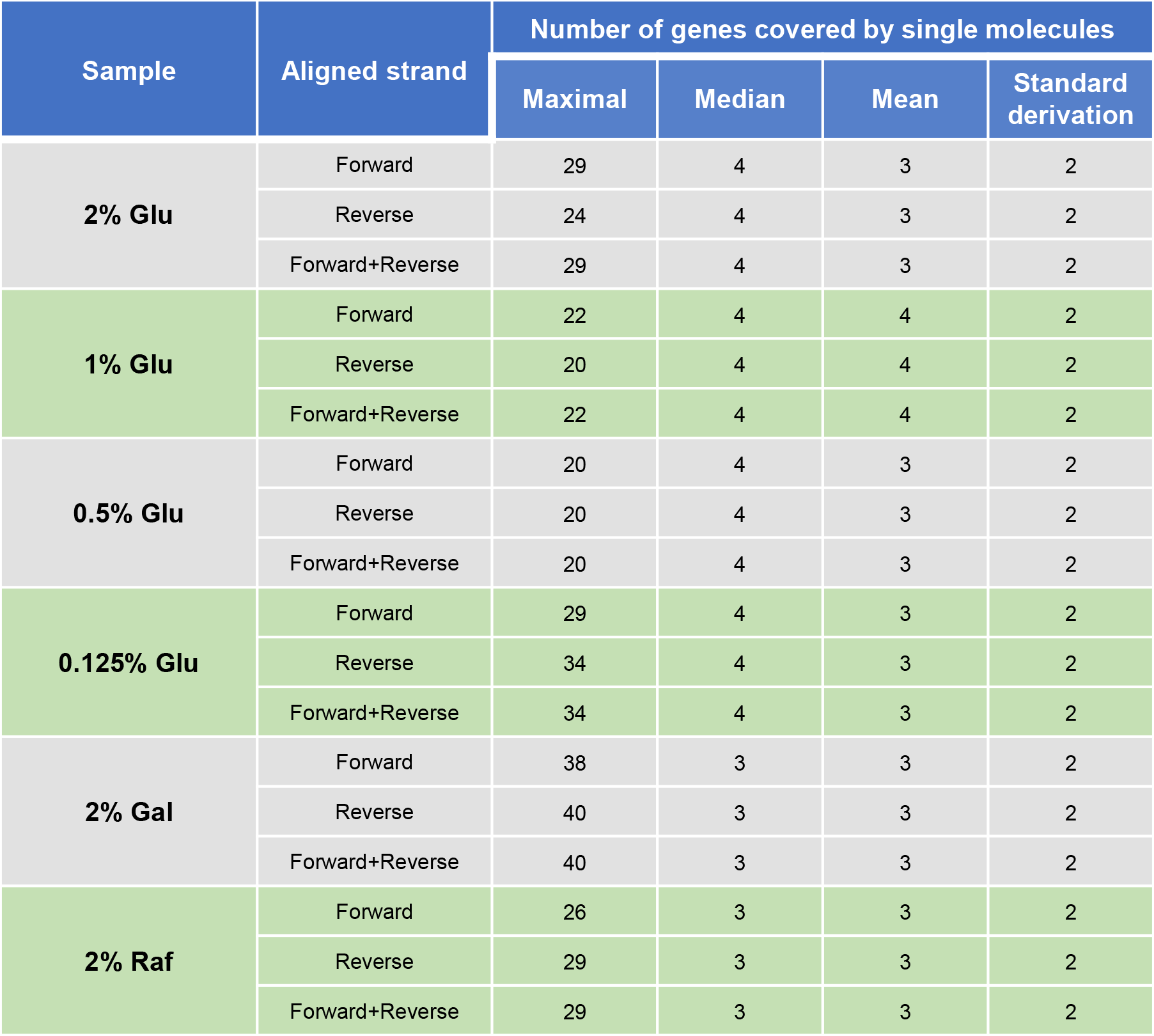
Number of genes covered by single molecules of MeSMLR-seq

**Table S4.** Statistics of biological samples and sequencing data used in this study

| Sample | Growth medium |  |  | Sequencing data |  |  |
| --- | --- | --- | --- | --- | --- | --- |
|  | Yeast extract | Peptone | Carbon source | MeSMLR-seq | Bulk-cell RNA-seq | Single-cell RNA-seq |
| 2% Glu | 1% | 2% | 2% Glucose | √ | √ | √ |
| 1% Glu | 1% | 2% | 1% Glucose + 1% Galactose | √ | √ |  |
| 0.5% Glu | 1% | 2% | 0.5% Glucose + 1.5% Galactose | √ | √ |  |
| 0.125% Glu | 1% | 2% | 0.125% Glucose + 1.875% Galactose | √ | √ |  |
| 2% Gal | 1% | 2% | 2% Galactose | √ | √ |  |
| 2% Raf | 1% | 2% | 2% Raffinose | √ | √ |  |

**Table S5.** Statistics of bulk-cell RNA-seq data

Statistics of bulk-cell RNA-seq data
| Sample | Biological replicate | Total number of read pairs | Alignment rate (%) |
| --- | --- | --- | --- |
| 2% Glu | Replicate 1 | 11462015 | 98.42 |
|  | Replicate 2 | 9091690 | 98.43 |
|  | Replicate 3 | 8459098 | 97.56 |
| 1% Glu | Replicate 1 | 9066796 | 98.38 |
|  | Replicate 2 | 10611557 | 98.29 |
|  | Replicate 3 | 10746015 | 98.32 |
| 0.5% Glu | Replicate 1 | 9691923 | 97.88 |
|  | Replicate 2 | 10111610 | 98.33 |
|  | Replicate 3 | 9994920 | 98.39 |
| 0.125% Glu | Replicate 1 | 11210531 | 98.15 |
|  | Replicate 2 | 9751364 | 98.20 |
|  | Replicate 3 | 9615422 | 97.70 |
| 2% Gal | Replicate 1 | 9614336 | 98.16 |
|  | Replicate 2 | 10500154 | 98.65 |
|  | Replicate 3 | 10784979 | 98.60 |
| 2% Raf | Replicate 1 | 10677473 | 98.70 |
|  | Replicate 2 | 10395431 | 98.62 |
|  | Replicate 3 | 9721105 | 98.50 |

**Table S6.**
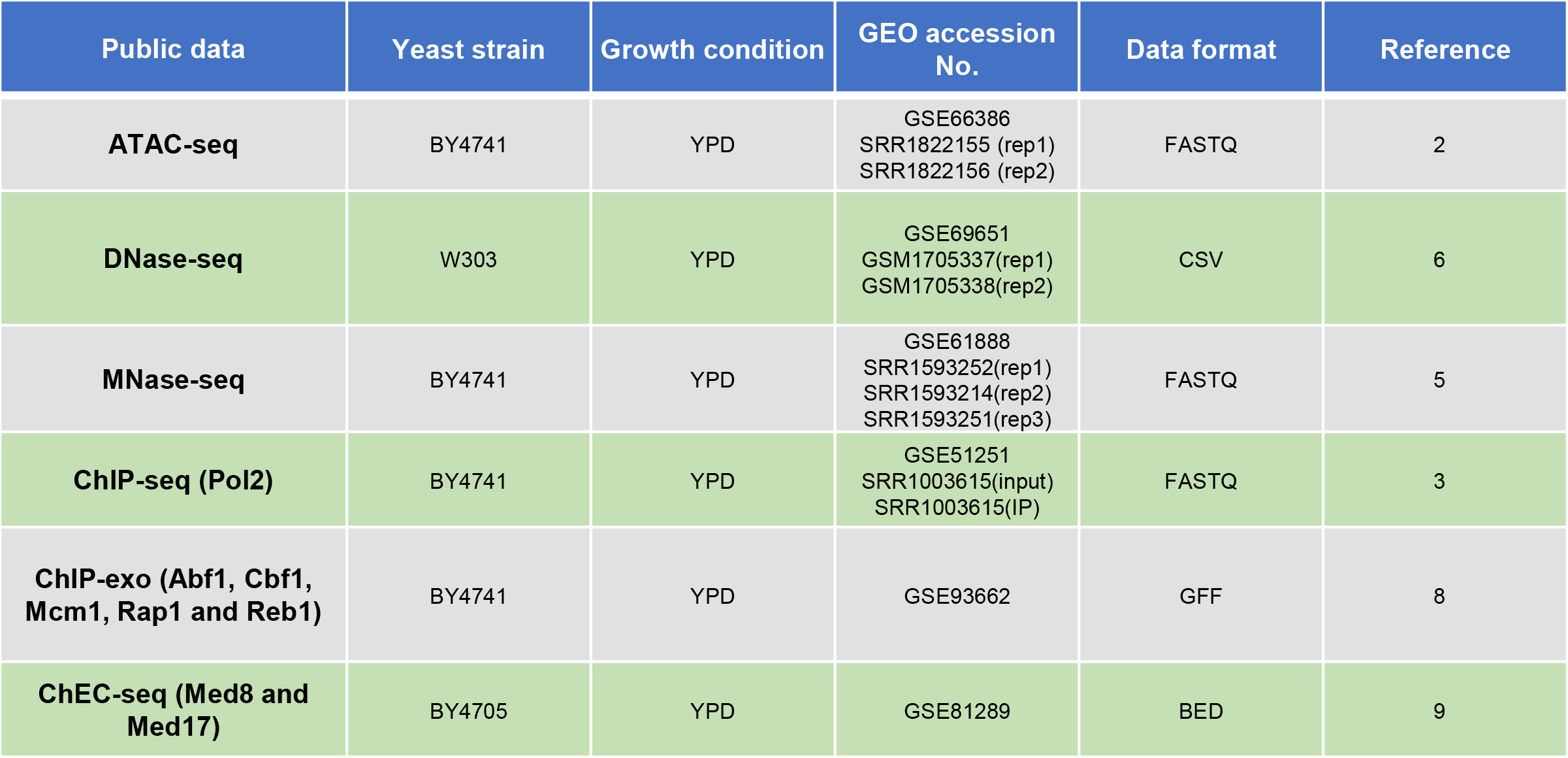
Statistics of public sequencing data used in this study

## REFERENCES

1. Luger K, Mader AW, Richmond RK, Sargent DF, & Richmond TJ (1997) Crystal structure of the nucleosome core particle at 2.8 A resolution. Nature 389(6648):251–260.

2. Bell O, Tiwari VK, Thoma NH, & Schubeler D (2011) Determinants and dynamics of genome accessibility. Nat Rev Genet 12(8):554–564.

3. Cui K & Zhao K (2012) Genome-wide approaches to determining nucleosome occupancy in metazoans using MNase-Seq. Methods Mol Biol 833:413–419.

4. Song L & Crawford GE (2010) DNase-seq: a high-resolution technique for mapping active gene regulatory elements across the genome from mammalian cells. Cold Spring Harb Protoc 2010(2):pdb prot5384.

5. Buenrostro JD, Wu B, Chang HY, & Greenleaf WJ (2015) ATAC-seq: A Method for Assaying Chromatin Accessibility Genome-Wide. Curr Protoc Mol Biol 109:21 29 21–29.

6. Kelly TK, et al. (2012) Genome-wide mapping of nucleosome positioning and DNA methylation within individual DNA molecules. Genome Res 22(12):2497–2506.

7. Ishii H, Kadonaga JT, & Ren B (2015) MPE-seq, a new method for the genome-wide analysis of chromatin structure. Proc Natl Acad Sci U S A 112(27):E3457–3465.

8. Wal M & Pugh BF (2012) Genome-wide mapping of nucleosome positions in yeast using high-resolution MNase ChIP-Seq. Methods Enzymol 513:233–250.

9. Bianco S, Rodrigue S, Murphy BD, & Gevry N (2015) Global Mapping of Open Chromatin Regulatory Elements by Formaldehyde-Assisted Isolation of Regulatory Elements Followed by Sequencing (FAIRE-seq). Methods Mol Biol 1334:261–272.

10. Buenrostro JD, et al. (2015) Single-cell chromatin accessibility reveals principles of regulatory variation. Nature 523(7561):486–490.

11. Jin W, et al. (2015) Genome-wide detection of DNase I hypersensitive sites in single cells and FFPE tissue samples. Nature 528(7580):142–146.

12. Lai B, et al. (2018) Principles of nucleosome organization revealed by single-cell micrococcal nuclease sequencing. Nature 562(7726):281–285.

13. Li L, et al. (2018) Single-cell multi-omics sequencing of human early embryos. Nat Cell Biol 20(7):847–858.

14. Clark SJ, et al. (2018) scNMT-seq enables joint profiling of chromatin accessibility DNA methylation and transcription in single cells. Nat Commun 9(1):781.

15. Pott S (2017) Simultaneous measurement of chromatin accessibility, DNA methylation, and nucleosome phasing in single cells. Elife 6.

16. Small EC, Xi L, Wang JP, Widom J, & Licht JD (2014) Single-cell nucleosome mapping reveals the molecular basis of gene expression heterogeneity. Proc Natl Acad Sci U S A 111(24):E2462–2471.

17. Rand AC, et al. (2017) Mapping DNA methylation with high-throughput nanopore sequencing. Nat Methods 14(4):411–413.

18. Simpson JT, et al. (2017) Detecting DNA cytosine methylation using nanopore sequencing. Nat Methods 14(4):407–410.

19. Payne A, Holmes N, Rakyan V, & Loose M (2018) BulkVis: a graphical viewer for Oxford nanopore bulk FAST5 files. Bioinformatics.

20. Capuano F, Mulleder M, Kok R, Blom HJ, & Ralser M (2014) Cytosine DNA methylation is found in Drosophila melanogaster but absent in Saccharomyces cerevisiae, Schizosaccharomyces pombe, and other yeast species. Anal Chem 86(8):3697–3702.

21. Needleman SB & Wunsch CD (1970) A general method applicable to the search for similarities in the amino acid sequence of two proteins. J Mol Biol 48(3):443–453.

22. Hughes AL & Rando OJ (2014) Mechanisms underlying nucleosome positioning in vivo. Annu Rev Biophys 43:41–63.

23. Weiner A, et al. (2015) High-resolution chromatin dynamics during a yeast stress response. Mol Cell 58(2):371–386.

24. Voss TC & Hager GL (2014) Dynamic regulation of transcriptional states by chromatin and transcription factors. Nat Rev Genet 15(2):69–81.

25. Schep AN, et al. (2015) Structured nucleosome fingerprints enable high-resolution mapping of chromatin architecture within regulatory regions. Genome Res 25(11):1757–1770.

26. Zhong J, et al. (2016) Mapping nucleosome positions using DNase-seq. Genome Res 26(3):351–364.

27. Yuan GC, et al. (2005) Genome-scale identification of nucleosome positions in S. cerevisiae. Science 309(5734):626–630.

28. Park D, Lee Y, Bhupindersingh G, & Iyer VR (2013) Widespread misinterpretable ChIP-seq bias in yeast. PLoS One 8(12):e83506.

29. Rossi MJ, Lai WKM, & Pugh BF (2018) Genome-wide determinants of sequence-specific DNA binding of general regulatory factors. Genome Res 28(4):497–508.

30. Grunberg S, Henikoff S, Hahn S, & Zentner GE (2016) Mediator binding to UASs is broadly uncoupled from transcription and cooperative with TFIID recruitment to promoters. EMBO J 35(22):2435–2446.

31. Paulo JA, O’Connell JD, Gaun A, & Gygi SP (2015) Proteome-wide quantitative multiplexed profiling of protein expression: carbon-source dependency in Saccharomyces cerevisiae. Mol Biol Cell 26(22):4063–4074.

32. Ozcan S & Johnston M (1999) Function and regulation of yeast hexose transporters. Microbiol Mol Biol Rev 63(3):554–569.

33. Jiang C & Pugh BF (2009) Nucleosome positioning and gene regulation: advances through genomics. Nat Rev Genet 10(3):161–172.

34. Li B, Carey M, & Workman JL (2007) The role of chromatin during transcription. Cell 128(4):707–719.

35. Petesch SJ & Lis JT (2008) Rapid, transcription-independent loss of nucleosomes over a large chromatin domain at Hsp70 loci. Cell 134(1):74–84.

36. Li G, Levitus M, Bustamante C, & Widom J (2005) Rapid spontaneous accessibility of nucleosomal DNA. Nat Struct Mol Biol 12(1):46–53.

37. Lipford JR & Bell SP (2001) Nucleosomes positioned by ORC facilitate the initiation of DNA replication. Mol Cell 7(1):21–30.

38. Cole HA, Howard BH, & Clark DJ (2011) The centromeric nucleosome of budding yeast is perfectly positioned and covers the entire centromere. Proc Natl Acad Sci U SA 108(31):12687–12692.

39. Dalal Y, Furuyama T, Vermaak D, & Henikoff S (2007) Structure, dynamics, and evolution of centromeric nucleosomes. Proc Natl Acad Sci U S A 104(41):15974–15981.

40. Schwartz S, Meshorer E, & Ast G (2009) Chromatin organization marks exon-intron structure. Nat Struct Mol Biol 16(9):990–995.

41. Tilgner H, et al. (2009) Nucleosome positioning as a determinant of exon recognition. Nat Struct Mol Biol 16(9):996–1001.

42. Lai WKM & Pugh BF (2017) Understanding nucleosome dynamics and their links to gene expression and DNA replication. Nat Rev Mol Cell Biol 18(9):548–562.

43. Rando OJ & Winston F (2012) Chromatin and transcription in yeast. Genetics 190(2):351–387.

44. Imielinski M, et al. (2019) Pore-C: using nanopore reads to delineate long-range interactions between genomic loci in the human genome. Available at: https://nanoporetech.com/resource-centre/pore-c-using-nanopore-reads-delineate-long-range-interactions-between-genomic-loci [Accessed Jan 23, 2019].

45. Li H & Durbin R (2010) Fast and accurate long-read alignment with Burrows-Wheeler transform. Bioinformatics 26(5):589–595.

46. Chen W, et al. (2014) Improved nucleosome-positioning algorithm iNPS for accurate nucleosome positioning from sequencing data. Nat Commun 5:4909.

47. Langmead B & Salzberg SL (2012) Fast gapped-read alignment with Bowtie 2. Nat Methods 9(4):357–359.

48. Zhang Y, et al. (2008) Model-based analysis of ChIP-Seq (MACS). Genome Biol 9(9):R137.

49. Boyle AP, Guinney J, Crawford GE, & Furey TS (2008) F-Seq: a feature density estimator for high-throughput sequence tags. Bioinformatics 24(21):2537–2538.

50. Huang da W, Sherman BT, & Lempicki RA (2009) Systematic and integrative analysis of large gene lists using DAVID bioinformatics resources. Nat Protoc 4(1):44–57.

## References for SI reference citations

1. Langmead B & Salzberg SL (2012) Fast gapped-read alignment with Bowtie 2. Nat Methods 9(4):357–359.

2. Schep AN, et al. (2015) Structured nucleosome fingerprints enable high-resolution mapping of chromatin architecture within regulatory regions. Genome Res 25(11):1757–1770.

3. Park D, Lee Y, Bhupindersingh G, & Iyer VR (2013) Widespread misinterpretable ChIP-seq bias in yeast. PLoS One 8(12):e83506.

4. Zhang Y, et al. (2008) Model-based analysis of ChIP-Seq (MACS). Genome Biol 9(9):R137.

5. Weiner A, et al. (2015) High-resolution chromatin dynamics during a yeast stress response. Mol Cell 58(2):371–386.

6. Zhong J, et al. (2016) Mapping nucleosome positions using DNase-seq. Genome Res 26(3):351–364.

7. Boyle AP, Guinney J, Crawford GE, & Furey TS (2008) F-Seq: a feature density estimator for high-throughput sequence tags. Bioinformatics 24(21):2537–2538.

8. Rossi MJ, Lai WKM, & Pugh BF (2018) Genome-wide determinants of sequence-specific DNA binding of general regulatory factors. Genome Res 28(4):497–508.

9. Grunberg S, Henikoff S, Hahn S, & Zentner GE (2016) Mediator binding to UASs is broadly uncoupled from transcription and cooperative with TFIID recruitment to promoters. EMBO J 35(22):2435–2446.

10. Stegle O, Teichmann SA, & Marioni JC (2015) Computational and analytical challenges in single-cell transcriptomics. Nat Rev Genet 16(3):133–145.

11. Anders S & Huber W (2010) Differential expression analysis for sequence count data. Genome Biol 11(10):R106.

12. Marcel M (2011) Cutadapt removes adapter sequences from high-throughput sequencing reads. EMBnet.journal 17(3).

13. Kim D, Langmead B, & Salzberg SL (2015) HISAT: a fast spliced aligner with low memory requirements. Nat Methods 12(4):357–360.

14. Trapnell C, et al. (2010) Transcript assembly and quantification by RNA-Seq reveals unannotated transcripts and isoform switching during cell differentiation. Nat Biotechnol 28(5):511–515.

